# Genetic and Morphological Insights Reveal the Hidden and Colorful Diversity in *Oscarella* Sponges

**DOI:** 10.1101/2024.10.26.620226

**Authors:** E. Guiollot, D. Guillemain, E. Renard, C. Borchiellini, Q. Schenkelaars

**Affiliations:** Aix-Marseille Université, CNRS, IRD, Avignon Université, IMBE, UMR 7263, Station Marine d′Endoume, Rue de la Batterie des Lions, 13007 Marseille, France; Marine Biology Department, Centre Scientifique de Monaco, Monaco, 98000, Monaco; Aix-Marseille Université, OSU Institut Pythéas, Campus de Luminy OCEANOMED, 163 avenue de Luminy, 13009 Marseille, France

**Keywords:** Porifera, Genetic markers, Species delimitation, Taxonomy, Biodiversity

## Abstract

The identification and classification of new taxa are crucial for understanding biodiversity. However, assigning samples to new taxa requires cautious and rigorous approaches. Historically, taxonomy has heavily relied on morphological traits, which can be subjective and may not always reflect underlying genetic divergence—particularly in organisms with few diagnostic morphological traits. A prime example is the sponge genus *Oscarella* (Homoscleromorpha), where species delimitation is challenged by the absence of spicules and limited morphological characters. Here, we address this gap by combining an extensive genetic dataset (192 specimens, five markers) with systematic photographic documentation. This approach enabled a robust assessment of *Oscarella* diversity in the Western Mediterranean, resulting in the identification of four species new to science. Multigene phylogenetic analyses also enabled us to propose an evolutionary scenario for color polymorphism. Moreover, our data highlighted critical limitations in current methodologies for studying *Oscarella*, including the low resolution of the widely used cytochrome c oxidase subunit I (cox1/COI) gene, the lack of genetic data for many species, and insufficient information on their geographical distribution. These issues, which mirror challenges across many taxa, highlight the urgent need for standardized genetic frameworks and comprehensive datasets to improve taxonomic resolution for each taxon.

## INTRODUCTION

Marine biodiversity remains largely unknown or underestimated in many ecosystems, despite their undeniable ecological, economic, and cultural importance^1^. Monitoring and implementing appropriate local management of these ecosystems therefore remain major challenges. One reason is that marine habitats are less accessible than most continental ones, requiring oceanographic vessels or diving equipment. Furthermore, they are often subject to strict regulations. As a result, marine ecosystems lag behind in their taxonomic description. Another factor contributing to this gap is the recurrent clustering of cryptic, non-vertebrate species under the names of presumably cosmopolitan taxa^2–5^. This issue stems from the frequent lack of clear morphological traits in numerous invertebrate groups, and from the fact that, although the use of molecular data for species description has gained wider acceptance, it is still not universally endorsed. Among the marine invertebrates facing such challenges, sponges are a prime example^6,7^.

Sponges (Porifera) are major components of many marine ecosystems. Together with other invertebrates such as corals and gorgonians (Cnidaria), they form structural frameworks that provide habitats for numerous organisms. Accordingly, many sponge species are considered ecosystem engineers, whose life cycles and persistence strongly influence the surrounding community. In the Mediterranean Sea, approximately 1,000 sponge species have been recorded, with a high level of endemism^8^. Among them, *Oscarella* species (Homoscleromorpha) play important roles, covering up to 37% of the substrate covered by sponges in some coralligenous ecosystems^9^. The first described *Oscarella* species was *O. lobularis* (Schmidt, 1862)^10^. Originally identified in the Mediterranean Sea, it has since been reported from various locations worldwide^11–30^, and is therefore often regarded as cosmopolitan. To date, 25 *Oscarella* species are formally recognized worldwide^31^ (**Table 1**), but their true diversity remains poorly understood, and their geographical distribution is still largely unresolved. In this context, the Bay of Marseille (Western Mediterranean Sea, France) can be considered the cradle of *Oscarella* research, as it is the area where species diversity has been most thoroughly documented, with seven species clearly identified (**Table 1**)^32,7,33^.

**Table 1:**
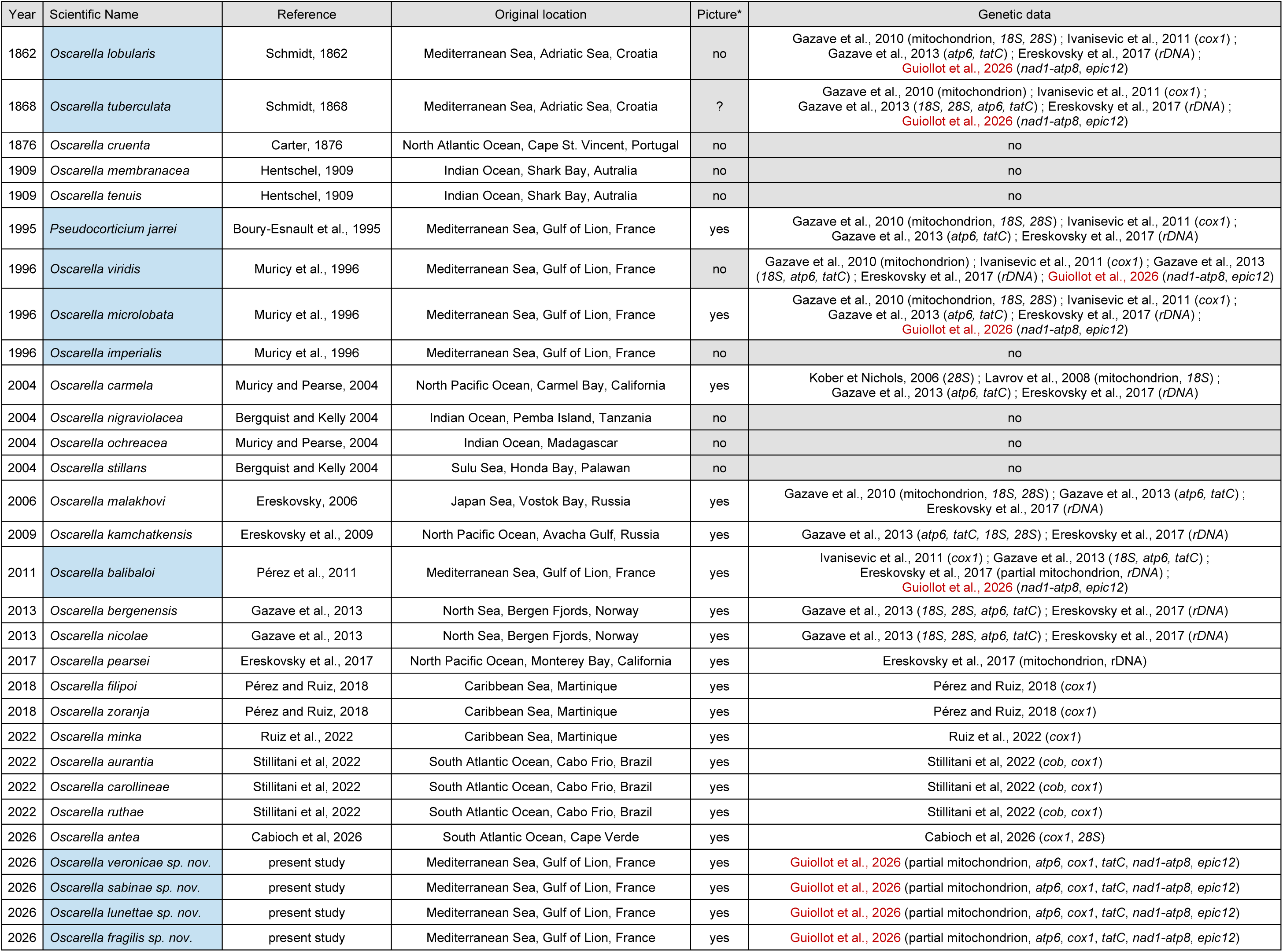
Chronological list of *Oscarella* accepted species according to Word Porifera Database. Species names underlined in blue indicate those found in the Bay of Marseille, Gulf of Lion, Western Mediterranean Sea. The table includes references for the original descriptions, along with the corresponding sampling locations for each species. Additionally, it provides information on whether informative images were included in the original descriptions and whether genetic data are available. The data presented in this study are highlighted in red. Note that, according to all phylogenetic studies, including this one, *Pseudocorticium jarrei* should be renamed *Oscarella jarrei*.

Species identification in sponges often relies on the description of their skeletons. The composition, shape, and size of small mineral structures called spicules, as well as the density and reticulation of organic fibers, vary greatly among species^34^. This approach, however, cannot be applied to *Oscarella*, as these sponges lack a skeleton^8^. Furthermore, *Oscarella* species generally share a relatively simple encrusting and lobate body forms with a limited number of conspicuous morphological characters available for species discrimination. Consequently, species identification in this genus has traditionally relied on few morphological characters (*e.g.*, lobe shape and size, and surface texture), cytological features (*e.g*., presence of cells containing paracrystalline inclusions), tissue consistency (fragile, soft, or cartilaginous), and color^7^. The limited number of traits, their low variability, and their inherent subjectivity make reliable species delimitation challenging. Since 2013, molecular analyses have therefore been systematically incorporated into the description of new species in this genus.

In the first genetic survey of *Oscarella*, two nuclear markers (*18S* and *28S ribosomal DNA*) and two mitochondrial markers (*ATP synthase membrane subunit 6*, *atp6*, and *twin-arginine translocase component*, *tatC*) were used to study a few samples collected worldwide^35^. While this approach was sufficient to discriminate most species and describe two new taxa, it failed to efficiently distinguish *O. lobularis* and *O. tuberculata*. In more recent studies, the *I3-M11* fragment of the widely used mitochondrial marker *cytochrome c oxidase subunit 1* (*cox1/COI*) was applied^36–40^. Unfortunately, it also failed to discriminate some closely related *Oscarella* species, such as *O. carollineae, O. filipoi* and *O. ruthae*^36^. These results may reflect (i) that the mentioned taxa are in fact single species, (ii) that the markers used are insufficiently informative to distinguish closely related species (*e.g*., due to low mutation rates or incomplete lineage sorting)^41^, or (iii) that species identification based on morphology and tissue consistency is not straightforward.

In this study, we collected 192 specimens of *Oscarella* spp. from eight locations across the Bay of Marseille to conduct a comprehensive genetic survey of local diversity. Our barcoding approach included (i) the three previously used mitochondrial protein-coding markers (*atp6*, *cox1*, and *tatC*), (ii) a newly developed mitochondrial marker covering the end of the NADH dehydrogenase subunit 1 gene, the beginning of the ATP synthase membrane subunit 8 gene, and the intergenic region (*nad1-atp8*), and (iii) a newly developed single-copy nuclear marker, an Exon Primed Intron Crossing (EPIC) locus (*epic12*)^42,43^. We compared these five markers and identified the most suitable for implementing a barcoding approach aimed at efficient species identification. In addition to species delimitation, partial mitochondrial genomes were obtained to clarify phylogenetic relationships and trace the evolution of color polymorphisms. Overall, this study represents the most comprehensive genetic investigation of this genus to date, both in terms of specimen sampling and the number of genetic markers.

## METHODS

### Specimen collection

We collected 192 *Oscarella* spp. specimens representing different color morphs from multiple locations across the Bay of Marseille (**Figure 1**). Each specimen was photographed (**Supplementary Figure 1**) and then assigned a unique identifier incorporating the sampling location and sample number (*e.g*., 3PP_01; **Supplementary Table 1**). Because our goal was to preserve the possibility of conducting complementary analyses on the same individuals (*e.g.,* immunohistochemistry, *in situ* hybridization) in addition to PCR amplification, we employed here a somewhat unconventional protocol previously developed in *Hydra*^44^. Accordingly, specimens were fixed overnight at 4 °C in a solution containing 4% paraformaldehyde (50% PBS-Tween 1‰, 25% seawater, 25% 16% paraformaldehyde aqueous solution). The following day, samples were washed in PBS-Tween, gradually transferred to increasing methanol concentrations, and finally stored in 100% methanol at −20 °C. When possible, one holotype and one paratype have been deposited at the Muséum National d’Histoire Naturelle de Paris for each new species. The corresponding accession numbers are provided in the Systematics section. All additional samples are maintained at our home institution (IMBE). For many specimens, small pieces of fresh tissue were also flash-frozen in liquid nitrogen and stored at −80 °C at the home institute (IMBE).

**Figure 1:**
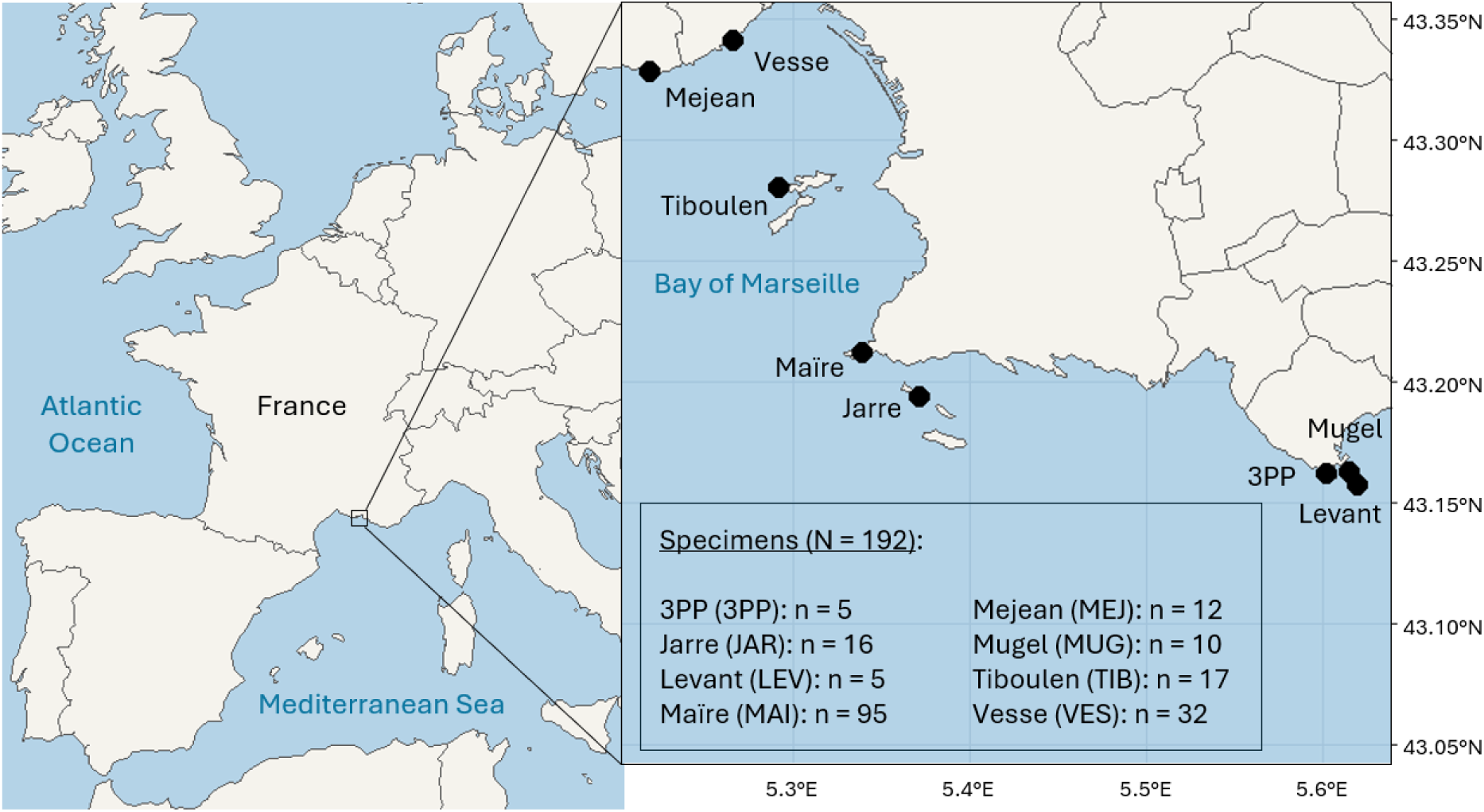
Sampling locations across the Bay of Marseille and number of samples at each location. Maps were generated in RStudio using the *sf*, *ggplot2*, and *rnaturalearth* packages. The high-resolution coastline basemap for the Bouches-du-Rhône department was obtained from IGN/Data.gouv.fr.

### Marker amplification, sequencing, and haplotype networks

In a previous study, we demonstrated that a direct-PCR approach based on rapid mechanical grinding with a pestle yields PCR products of quality comparable to standard extraction-based methods, while requiring substantially less time and reducing costs^44^. For this reason, no DNA extraction was performed in the present study. Instead, a small piece of fixed tissue (~3 mm^2^) was ground in 1 mL of water with a pestle to serve as crude template. Mitochondrial (*atp6*, *cox1*, *nad1-atp8*, *tatC*) and nuclear (*epic12*) markers were amplified using GoTaq® G2 Flexi DNA polymerase (Promega). PCR reactions were performed in 20 µL volumes containing 1X buffer, 0.2 mM dNTPs, 1 µM primers, 1.5 mM MgCl₂, 0.5 U GoTaq, and 3 µL of template. Amplifications were carried out for 30 cycles. Primer sequences, amplicon sizes, and PCR annealing temperatures are listed in **Table 2**. PCR products were verified on 1% agarose gels and submitted to Eurofins Genomics for Sanger sequencing. All sequences generated in this study have been deposited in the NCBI database (**Supplementary Table 2**). Sequences of *nad1-atp8* and *tatC* were aligned using MUSCLE^45^, whereas other markers did not require any alignment because the sequences are highly conserved, identical in length, and contain no indels. For these markers, we simply trimmed the sequence ends to standardize their length. Haplotype networks were constructed using the Minimum Spanning Network function in PopArt^46^.

**Table 2:**
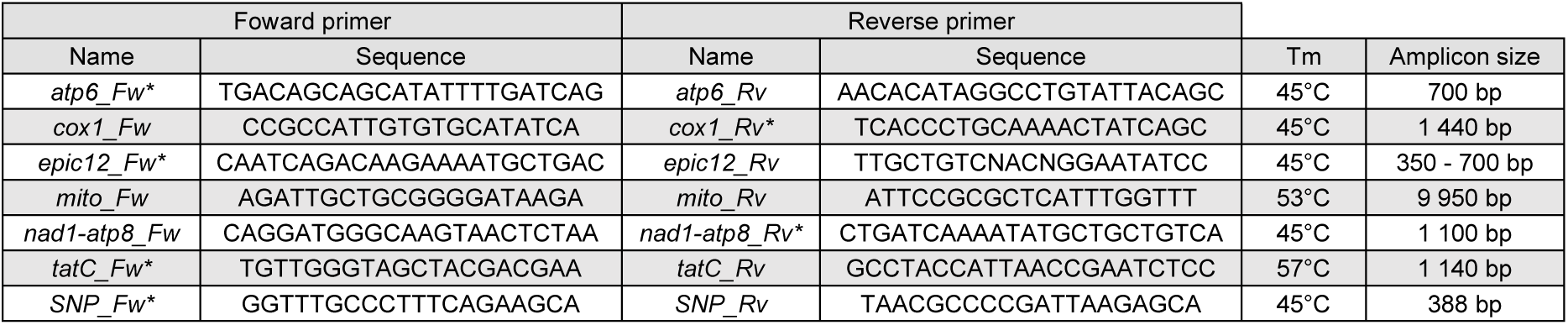
Information on PCR primers designed for the genus *Oscarella*, including primer sequences, Tm, and amplicon size. (*) indicates primers used for Sanger sequencing.

### Mitochondrial genome amplification of new species, sequencing, and phylogenetic reconstruction

DNA was extracted from 25 mg of flash-frozen tissue using the DNeasy® Blood & Tissue Kit (Qiagen). Long PCRs were performed with GoTaq® Long PCR Master Mix (Promega) containing 1X mix, 1 µM primers, and 5 µg DNA. Primer sequences, amplicon sizes, and melting temperatures are provided in **Table 2**. PCR amplicons were visualized on 0.5% agarose gels, purified using the Qiagen DNA extraction kit, and sequenced via Oxford Nanopore Technologies services at Eurofins Genomics. Partial mitochondrial genomes have been deposited in the NCBI database (**Supplementary Table 2**). Amplicons encompassed eight mitochondrial protein-coding genes (*cox1*, *nad1*, *atp8*, *atp6*, *cox3*, *atp9*, *nad4*, and *nad6*), which were concatenated for each species. Resulting sequences were aligned with MUSCLE^45^, and poorly aligned positions were removed using Gblocks with the following settings: minimal conservation for a flank position 85%, maximal contiguous non-conserved positions 8 bp, minimal block length 10 bp, and no gaps allowed in final blocks^47^, retaining 98% of the original 6591 positions. The best-fit nucleotide substitution model was estimated using jModelTest^48^. Phylogenetic reconstruction was performed in MEGA^49^ using the Maximum Likelihood method with the General Time Reversible model, a gamma-distributed substitution rate (four categories), and a proportion of invariable sites (GTR + I + G). Branch support was assessed with 1000 bootstrap replicates.

## RESULTS

### Comprehensive genetic survey of *Oscarella* species in the Bay of Marseille

PCR amplification of the *atp6* marker using crude macerates as templates was successful for the majority of specimens (95% amplification success). Consequently, *atp6* haplotypes were used as a practical initial partitioning tool, at a stage where morphological identification was not yet possible. Analysis of *atp6* sequences from 183 newly collected specimens identified 11 distinct *atp6* haplotypes, labeled a to k (**Figure 2A**). Comparisons with NCBI reference mitochondrial genomes assigned *atp6* haplotypes a and d to *O. tuberculata*, b to *O. lobularis*, g to *O. viridis*, and h to *O. microlobata*. In contrast, none of the *atp6* sequences matched *O. balibaloi* or *P. jarrei*, so we incorporated available mitochondrial genome sequences for these two species into the *atp6* network (*atp6* haplotypes l and m, respectively). Species identification could not be determined for the remaining six *atp6* haplotypes. Notably, *atp6* haplotypes e and j (corresponding to ten and one samples, respectively) branched directly on the main *O. tuberculata* haplotype (*atp6* haplotype a) without additional connections (**Figure 2A**).

**Figure 2:**
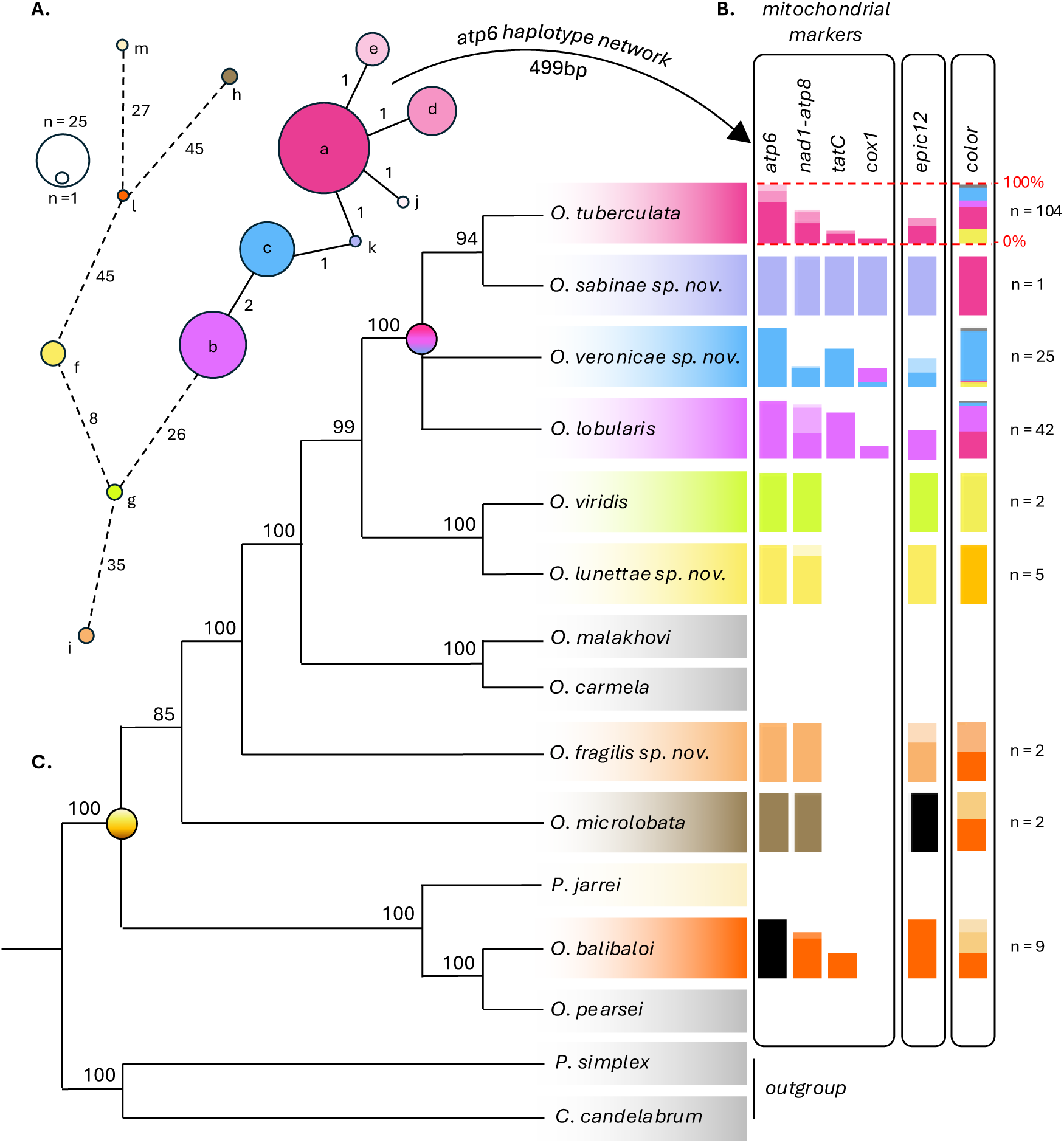
Species Delimitation Analysis of *Oscarella spp.* mapped onto a Maximum Likelihood (ML) Phylogenetic Tree. (A) Haplotype network based on the *atp6* marker. Each haplotype (a, b, c, etc.) is depicted as a circle, with the size of the circle proportional to the number of specimens represented. Numbers between haplotypes indicate the number of mutations separating them. (B) Schematic representation of species partitions derived from single-locus sequences, either mitochondrial (*atp6*, *nad1-atp8*, etc.) or nuclear (*epic12*). The proportion of each color morphs within each species is also provided (color). For instance, among the 104 *O. tuberculata* specimens collected, 39% were pink, 22% yellow, 20% blue, 9% purple, 9% black, and 1% beige. Detailed specimen information is provided in **Supplementary Table 1** and haplotype networks are shown in **Supplementary Figure 2**. (C) Maximum likelihood (ML) phylogenetic cladogram of *Oscarella* species based on eight mitochondrial protein-coding genes (*cox1*, *nad1*, *atp8*, *atp6*, *cox3*, *atp9*, *nad4*, and *nad6*). Only nodes with bootstrap values greater than 85% (1000 BS replicates) are shown. Dots on the tree indicate the inferred ancestral color state of the common ancestor of *Oscarella* and the last common ancestor of *O. lobularis*, *O. sabinae* sp. nov., *O. tuberculata*, and *O. veronicae* sp. nov. Note that *Pseudocorticium jarrei* is proposed to be renamed *Oscarella jarrei,* in agreement with the present results and previous ones (**Table 1**).

To (i) validate the *atp6* data, (ii) assess whether the six unresolved *atp6* haplotypes represent new taxa, and (iii) obtain genetic information for the nine specimens where *atp6* amplification failed, we sequenced three additional mitochondrial markers in a subset of individuals: *nad1-atp8* (66% of specimens), *tatC* (40%), and *cox1* (14%) (**Supplementary Figure 2**). Overall, the partitions obtained from these markers—except for *cox1*—were either fully consistent or compatible with the *atp6* data (**Figure 2B**, **Supplementary Table 1**, **Supplementary Figure 2**). For instance, *atp6* haplotype b and *tatC* haplotype a were consistently associated, with all individuals carrying one also carrying the other (*i.e.* consistent). Alternatively, only individuals with *atp6* haplotype c carried *nad1–atp8* haplotype e or m, and all individuals with *nad1–atp8* haplotype e or m carried *atp6* haplotype c (*i.e.* compatible). Specimens for which *atp6* amplification failed were identified as *O. balibaloi* using *nad1-atp8* and *tatC*, while *atp6* haplotypes a, d, e, and j must be grouped together as *O. tuberculata* according to *nad1-atp8* and *tatC* partitioning. These clustering patterns were further corroborated by the newly developed nuclear marker *epic12*, which yielded data consistent or compatible with the *atp6* haplotype network (**Figure 2B**, **Supplementary Table 1**, **Supplementary Figure 2**). Amplification of *epic12* failed for *O. microlobata*.

Combined analyses of *atp6*, *nad1-atp8, tatC* and *epic12* markers strongly suggest that individuals showing different *atp6* haplotypes correspond to distinct lineages, likely representing separate species. Indeed, none of these lineages share alleles for the mitochondrial markers (*atp6*, *nad1–atp8*, *tatC*) nor for the nuclear marker (*epic12*), despite their sympatric distribution within the Bay of Marseille (**Supplementary Table 1**). Importantly, the nuclear dataset includes a substantial sample size (*n* = 103 individuals), providing robust support for the absence of allele sharing. This pattern strongly suggests that no ongoing gene flow occurs among these lineages, which is consistent with a classical biological definition of species. Thus, at least ten *Oscarella* species inhabit the bay, including four new to science. We propose the following names: *O. veronicae* sp. nov. (*atp6* haplotype c), *O. sabinae* sp. nov. (*atp6* haplotype k), *O. fragilis* sp. nov. (*atp6* haplotype i), and *O. lunettae* sp. nov. (*atp6* haplotype f). Formal descriptions with morphological and genetic diagnosis are provided in the *Systematics* section (**Figure 3 and 4**).

**Figure 3:**
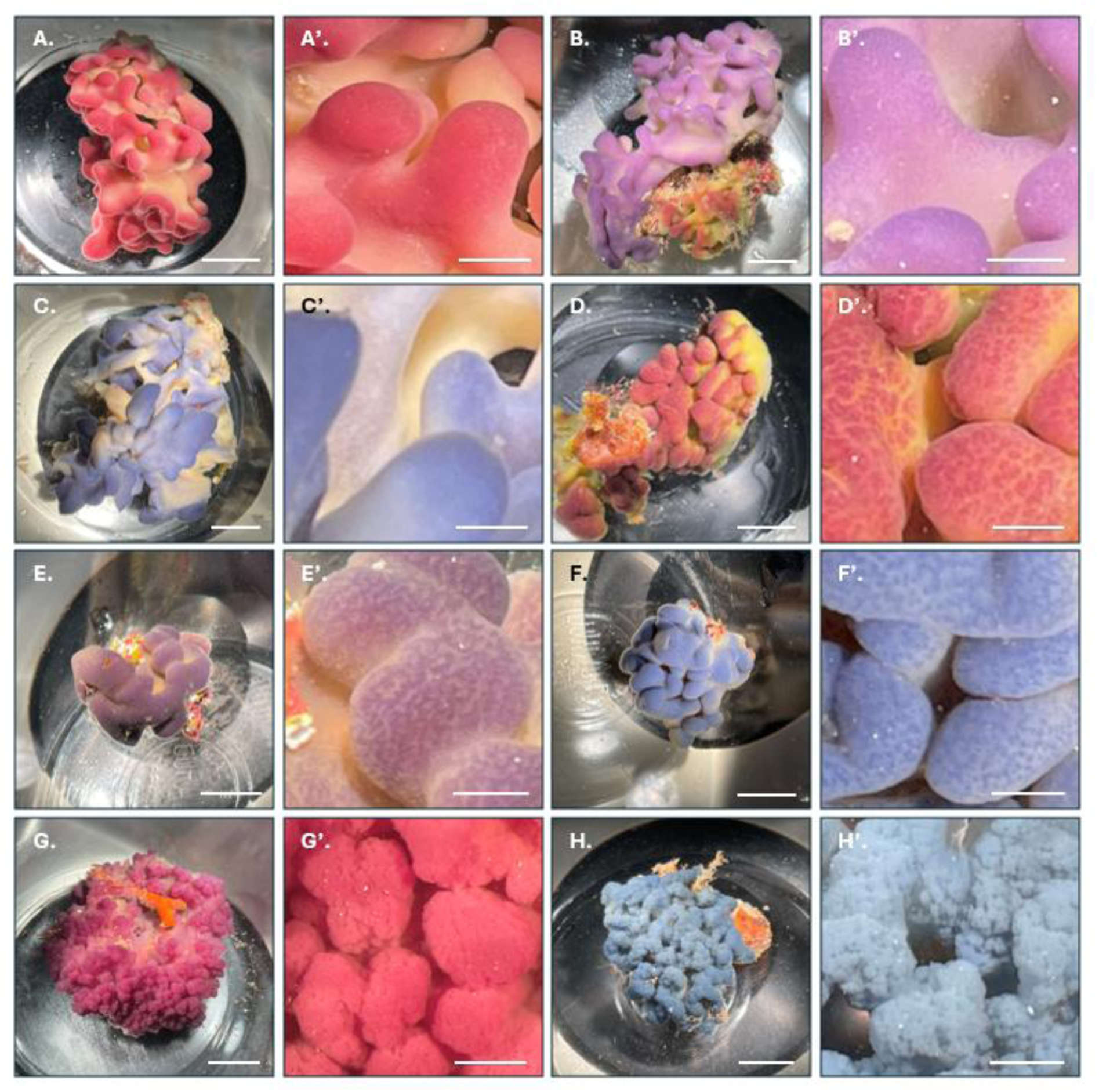
Comparative external morphology of *Oscarella* species within the polychromatic complex. (A to C’) *Oscarella lobularis.* (D to F’) *Oscarella tuberculata.* (G and G’) *Oscarella sabinae* sp. nov. (H and H’) *Oscarella veronicae* sp. nov. (A, B, C, D, E, F, G and H) Top-view photographs of the entire specimens. Scale: 1 cm. (A’, B’, C’, D’, E’, F’, G’ and H’) Close-up view of the lobes. Scale: 0.25 cm.

**Figure 4:**
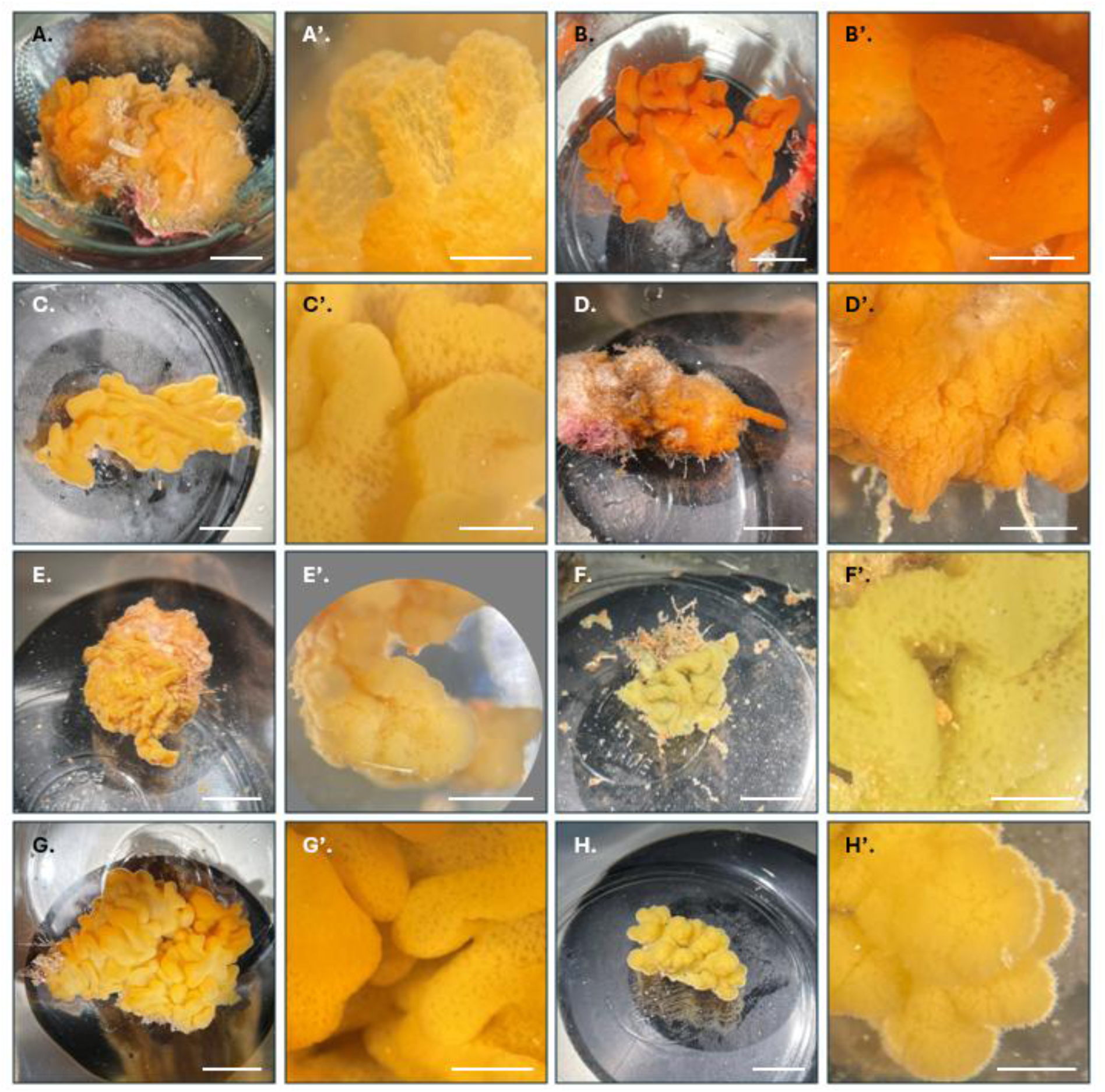
Comparative external morphology of yellow and orange *Oscarella* specimens. (A and A’) *Oscarella fragilis* sp. nov. (B to C’) *Oscarella balibaloi*. (D to E’) *Oscarella microlobata*. (F and F’) *Oscarella viridis*. (G and G’) *Oscarella lunettae*. (H and H’) *Oscarella veronicae* sp. nov. The somewhat hispid appearance corresponds to a particular physiological state of the specimen associated with the early stage of a budding event and should not be considered a characteristic feature of *O. veronicae*. (A, B, C, D, E, F, G and H) Top-view photographs of the entire specimens. Scale: 1 cm. (A’, B’, C’, D’, E’, F’, G’ and H’) Close-up view of the lobes. Scale: 0.25 cm.

### Phylogenetic relationships reveal a complex of four polychromatic species

Haplotype network analyses indicated that four *Oscarella* species are closely related. Specifically, *O. tuberculata* and *O. lobularis* consistently clustered with *O. sabinae* sp. nov. and *O. veronicae* sp. nov. (**Figure 2A**, **Supplementary Figure 2**). Regardless of the marker used (*atp6, nad1-atp8, tatC, cox1,* or *epic12*), genetic distances among haplotypes within this complex were very short compared to the distances between this complex and the nearest neighboring species (*e.g*., 0.4% vs. 5.2% for *atp6*). Similarly, *O. viridis* and *O. lunettae* sp. nov. were closely related across all markers (*e.g.*, 1.6% for *atp6*).

To further resolve relationships among *Oscarella* species, we sequenced 9,874 bp of the mitochondrial genome for one specimen of each new species. The resulting phylogenetic tree strongly supported (BS = 100%) the close relationship of *O. tuberculata*, *O. lobularis*, *O. sabinae* sp. nov., and *O. veronicae* sp. nov. (**Figure 2C**, **Supplementary Figure 3**). The analysis also confirmed that *O. lunettae* sp. nov. is the sister species to *O. viridis* (BS = 100%), whereas *O. fragilis* sp. nov. formed a distinct lineage, diverging before the *O. carmela* / *O. malakovi* clade, two North Pacific species (BS = 100%).

Given the importance of color for species identification in *Oscarella*, we examined color polymorphism in each species (**Figure 2B, Supplementary Figure 1**). *O. tuberculata*, *O. lobularis*, and *O. veronicae* sp. nov. exhibited a broad color range, including pink and blue morphotypes (**Figure 3**). Notably, the only *O. sabinae* sp. nov. specimen was bright pink, whereas other *Oscarella* species displayed colors from white to bright or rusty orange, including shades of light brown, beige, and yellow (**Figure 4**). Polychromatic species with pink, violet/purple, or blue morphotypes formed a monophyletic group. Consequently, color is a reliable criterion for distinguishing species within the complex from those outside it (although yellow specimens also occur within the complex), but it is not sufficient on its own for definitive species identification.

## DISCUSSION

### Towards a standardized approach for species identification in the *Oscarella*

By the late 1990s and early 2000s, PCR and sequencing were established as reliable tools for species identification and discovery. In *Oscarella*, barcoding was first applied only in 2013, using mitochondrial markers (*atp6* and *tatC*) and standard nuclear markers (18S, 28S)^35^. Despite some discrepancies, both mitochondrial markers showed overall more consistent results with the *a priori* assignment of individuals to putative species than the two nuclear markers. This may be due to ribosomal genes existing in multiple tandem copies, and while they are expected to evolve in a concerted manner, slightly divergent paralogues can persist within the same genome^50,51^. Amplification of these mixed copies can thus complicate phylogenetic interpretation. Consequently, subsequent studies solely relied on alternative mitochondrial markers such as *cox1 I3–M11* and *cob*^36–38,52^ (**Table 1**). Nevertheless, *cox1* failed to separate *O. lobularis* from *O. veronicae sp. nov.* in our study, as observed in other recent studies for *O. carollineae*, *O. filipoi*, and *O. ruthae*^36^. This highlights the broader limitations of *cox1/COI* for species delimitation in some taxa^53,54^. In the *Oscarella* context, *atp6*, *nad1-atp8*, and *tatC* outperformed *cox1* in discriminating closely related species, even though we observed high *cox1* amplification success rates. Furthermore, to bypass issues observed with standard nuclear markers (28S and 18S), we developed the novel single-copy nuclear marker *epic12*^42,43^, which can be reliably amplified across relatively distant *Oscarella* species. Interestingly, this new nuclear marker provided results strongly consistent with the mitochondrial markers. Accordingly, for reliable cross-study comparisons and robust phylogenetic reconstruction, we recommend that future species descriptions include partial mitochondrial genomes encompassing all relevant protein-coding markers, the nuclear marker *epic12*, and photographic documentation. Achieving a uniform dataset will also require resampling species for which genetic data and/or images are currently unavailable/incomplete, as illustrated by *O. imperialis* in the Western Mediterranean Sea, and by many others worldwide (**Table 1**), which cannot currently be identified with confidence^55,56^.

### Expanding geographical knowledge beyond type localities

A major challenge in understanding *Oscarella* diversity is the poorly known distribution of each species. For most of them, records are limited to their type localities. Exceptions such as *O. lobularis* and *O. tuberculata*, first described in the Adriatic Sea (Croatia)^10,57^, exhibit broader Mediterranean distributions and have even been reported from the North Atlantic Ocean^11–30,58–62^. However, most of these records lack supporting evidence (no genetic data and few photographs) and should therefore be considered potential sightings rather than confirmed occurrences. Comprehensive genetic sampling across the Mediterranean and beyond is essential to accurately map species distributions, inform conservation efforts (endemic *vs*. cosmopolitan species), and reveal new taxa. Our study exemplifies this approach, identifying four new species in the Bay of Marseille and increasing local *Oscarella* species richness by 57%. This is particularly relevant for the Homoscleromorpha, which comprise only 136 accepted species globally (<1.4% of total sponge diversity)^11^.

### Evolutionary insights and biochemical mysteries of color polymorphysm

Our results provide strong evidence that color polymorphism in *Oscarella* has a clear evolutionary signal. *O. tuberculata* and *O. lobularis* are truly polychromatic species, with morphotypes ranging from pink to blue and violet, whereas other formally described species examined here exhibit more subdued hues such as cream, yellow, orange, or brown^7,32,33,35–38,63–66^. Phylogenetic analyses including the new species suggest that remarkable color polymorphism arose within a single lineage uniting *O. tuberculata*, *O. lobularis*, *O. veronicae* sp. nov., and *O. sabinae* sp. nov., and thus represents a synapomorphy of this clade. By contrast, the orange *O. fragilis* sp. nov. and the yellow *O. lunettae* sp. nov. fall outside this group, following the broader pattern. Exploring additional species such as the Brazilian *O. ruthae* and *O. filipoi*, which also show signs of color polymorphism (although no blue morphs have been observed)^36^, could shed further light on whether this trait evolved once or multiple times within the genus.

Despite the dazzling colors displayed by many sponges, the biochemical basis of their pigmentation remains a largely uncharted territory. To date, most identified pigments are conventional specialized metabolites—quinones, alkaloids, terpenes, or carotenoids—responsible for the more common yellow, red, or black hues^67^. In contrast, true blue pigments are exceptionally rare across the tree of life, with coloration in most animals arising instead from structural effects such as nanocrystals or microscopic scales^68^. Only a handful of genuine blue pigments have been identified so far, including anthocyanins in plants^69^, GFP-like chromoproteins in cnidarians^70,71^, and tunichrome peptides in tunicates^72^. The presence of vivid blue morphotypes in *Oscarella* therefore raises a fascinating question. Deciphering the molecular basis of this polymorphism could open new perspectives in both evolutionary biology and natural product chemistry.

## SYSTEMATICS

Phylum: Porifera Grant, 1836.

Class: Homoscleromorpha Bergquist, 1978. Order: Homosclerophorida Dendy, 1905.

Family: Oscarellidae Lendenfeld, 1887. Genus: *Oscarella* Vosmaer, 1884.

### *Oscarella fragilis* sp. nov. (Schenkelaars & Guiollot)

#### Holotype

IMBE-A059-JAR01: Collected from France, Mediterranean Sea, Marseille, Jarre (43.19°N, 5.36°W), at a depth of 14-17 m by Dorian Guillemain on 2023-04-26 (JAR_01, **Figure 4A and A’**). Specimen fixed in 4% paraformaldehyde and preserved in absolute methanol.

#### Paratype

IMBE-A144-TIB11: Collected from France, Mediterranean Sea, Marseille, Tiboulen du Frioul (43.2799°N, 5.2860°W), at a depth of 30-35 m by Dorian Guillemain on 2023-06-28 (TIB_11, **Supplementary Figure 1**). Specimen fixed in 4% paraformaldehyde and preserved in absolute methanol.

#### Molecular diagnosis

Molecular analysis provides strong evidence for recognizing *O. fragilis* sp. nov. as a distinct species, with its genetic sequences showing high divergence from all other known *Oscarella* species (**Figure 2, Supplementary Figure 3**). The most striking example of this is the *epic12* sequences, which do not align with any haplotypes found in other species (*epic12* haplotype i and p, **Supplementary Figure 2**).

#### Morphological diagnosis

Although the entire surface of this species is lobed, the lobes form transparent, very membranous lamellae, a feature that starkly contrasts with those of other species in the genus (**Figure 3** and **4**). Furthermore, the tissue is more fragile than in *O. viridis* yet considered delicate^7^. While *O. viridis* tends to crumble, *O. fragilis* sp. nov exhibits a soft, almost mucous-like consistency, making manipulation with forceps particularly challenging (**Table 3**).

**Table 3:** Main morphological, anatomical, and cytological characters used for the discrimination of *Oscarella* species recorded in the Bay of Marseille. All species considered here lack a mineral skeleton, and the aquiferous system is characterized by a sylleibid canal system, eurypylous choanocyte chambers, and a basal cavity. Ectosome thickness and choanocyte chamber size also show broadly overlapping ranges among most species. Although these characters are important for documenting and describing *Oscarella* species, our comparison indicates that their discriminatory power is limited, particularly among closely related species. (*) Data for this species is based on observations from a single specimen. (**) Data provided are representative of most specimens but may show some variation. (***) Values marked “max.” correspond to the largest choanocyte chambers observed in the sections examined in the present study. (n.a.) Indicates that no data was obtained because the tissue samples were of insufficient quality.

|  | Reference | Skeleton | Shape | Surface | Colour | Consistency | Ectosome thickness | Chamber size*** |
| --- | --- | --- | --- | --- | --- | --- | --- | --- |
| *O. tuberculata* | Murcy et al., 1996 (7) | absent | Thick lobate | Wrinkled | Variable | Cartilaginous | 5 - 90 | 40 - 75 |
|  | Guiollot et al., 2026 (present) | absent | Broad, thick, elongated lobes | Smooth | Gradient, highly variable, often mottled | Cartilaginous | 35 - 93 | 52 - 65 (max.) |
| *O. sabinae* * | Guiollot et al., 2026 (present) | absent | Thick, rounded lobes | Verrucose (cauliflower-like) | Pink | Soft | 6 - 61 | 56 - 68 (max.) |
| *O. veronicae* | Guiollot et al., 2026 (present) | absent | Thick, rounded lobes | Verrucose (cauliflower-like)** | Blue, sometimes pink or yellow | Variable | 14 - 58 | 73 - 99 (max.) |
| *O. lobularis* | Murcy et al., 1996 (7) | absent | Thick lobate | Smooth | Variable | Soft | 5 - 50 | 35 - 90 |
|  | Guiollot et al., 2026 (present) | absent | Broad, thick, elongated lobes | Smooth | Gradient, pink, purple or blue | Soft | 7 - 48 | 63 - 74 (max.) |
| *O. viridis* | Murcy et al., 1996 (7) | absent | Thin lobate | Rugose | Light green | Very soft, fragile | 10 - 15 | 30 - 75 |
|  | Guiollot et al., 2026 (present) | absent | Broad, thin, elongated lobes | Rugose (ostia easily visible) | Yellow, light green | Fragile | n.a. | n.a. |
| *O. lunettae* | Guiollot et al., 2026 (present) | absent | Broad, thick, elongated lobes | Rugose (ostia easily visible) | Gradient, orange to yellow | Soft | 4 - 40 | 69 - 85 (max.) |
| *O. fragilis* | Guiollot et al., 2026 (present) | absent | Broad, thick, elongated lobes | Membranous | Orange | Fragile | n.a. | 62 - 73 (max.) |
| *O. microlobata* | Murcy et al., 1996 (7) | absent | Thin lobate | Rugose | Light brown | Soft, Fragile | 10 - 50 | 40 - 75 |
|  | Guiollot et al., 2026 (present) | absent | Thin, rounded lobes | Verrucose | Light brown, yellow, or orange | Soft | 9 - 44 | 49 - 75 (max.) |
| *O. balibaloï* | Guiollot et al., 2026 (present) | absent | Broad, thick, elongated lobes | Rugose (ostia easily visible) | Light brown, yellow, or orange | Soft | 7 - 47 | 67 - 90 (max.) |

#### Morphological description

Like other *Oscarella* species, *O. fragilis* sp. nov. is an encrusting sponge characterized by its irregular growth forms with broad, thick and elongated lobes (**Figure 4B to C’**, **Table 3**). Specimen covers areas of 15–20 cm^2^ with a thickness of 15–25 mm. The specimens collected in this study exhibit distinct colorations. One is pale orange (JAR_01, **Figure 4A**), while the other is bright orange (TIB_11, **Supplementary Figure 1**). This color pattern is reminiscent of *O. balibaloi* (*e.g.* 3PP_03 and MEJ_11, **Figure 4B to C’**, **Table 3**) and *O. microlobata* (*e.g.* JAR_14 and JAR_11, **Figure 4D to E’**, **Table 3**).

#### Cytological description

This species is very fragile; consequently, most of the ectosome was lost during specimen processing (sampling, fixation, and/or sectioning), preventing us from assessing its thickness. The aquiferous system appears similar to that described in other *Oscarella* species (**Supplementary Figure 4**). The largest choanocyte chambers observed in the sections measured 62–73 µm in diameter, (**Table 3**).

#### Ecology and distribution

One specimen was found encrusting red algae, while the other was encrusted on another organism, possibly the bryozoan *Smittina cervicornis* (TIB_11, **Supplementary Figure 1**). The first specimen was collected at a depth of 14–17 m, while the second was found at 30–35 m. Given the limited data, it is currently premature to draw definitive conclusions about the ecology and distribution of this species.

### *Oscarella lunettae* sp. nov. (Schenkelaars & Guiollot)

#### Holotype

*Muséum National d’Histoire Naturelle de Paris*, MNHN-IP-2019-3414, IMBE-A205-LEV04: Collected from France, Mediterranean Sea, Marseille, Levant (43.09288°N, 5.37365°W), at a depth of 40 m by Dorian Guillemain on 2024-03-07 (LEV_04, **Figure 4G and G’**). Specimen fixed in 4% paraformaldehyde and preserved in absolute methanol.

#### Paratype

*Muséum National d’Histoire Naturelle de Paris*, MNHN-IP-2019-3415, IMBE-A206-LEV05: Collected from France, Mediterranean Sea, Marseille, Levant (43.09288°N, 5.37365°W), at a depth of 40 m Dorian Guillemain on 2024-03-07 (LEV_05, **Supplementary Figure 1**). Specimen fixed in 4% paraformaldehyde and preserved in absolute methanol.

#### Other material examined

Station Marine d’Endoume, IMBE: three additional specimens collected from Levant (43.09288°N, 5.37365°W) and Tiboulen du Frioul (43.2799°N, 5.2860°W) (**Supplementary Table 1** and **Supplementary Figure 1**). Specimen fixed in 4% paraformaldehyde and preserved in absolute methanol.

#### Molecular diagnosis

Molecular data strongly support the recognition of *O. lunettae* sp. nov. as a distinct species, as its sequences show significant divergence from all currently available *Oscarella* data (**Figure 2**, **Supplementary Figure 3**), even from the closely related species, *O. viridis*, from which it differs by 118 substitutions across the 9,963 bp of mitochondrial sequence available for comparison (see **Supplementary Table 3** for ONT sequencing quality and coverage). Although the *epic12* sequence in *O. lunettae* has the same length as in most other species (51 bp), it does not align with them. In fact, *epic12* haplotype g (**Supplementary Figure 2**) does not share more than three consecutive residues with any same-length sequence. Instead, it more closely resembles *epic12* haplotype j (corresponding to *O. viridis*), which is longer (55 bp). Yet, the two sequences remain clearly distinct, differing by 13 diagnostic residues.

#### Morphological diagnosis

Although this species shares beige and orange tones with *O. balibaloi*, *O. microlobata*, and *O. fragilis* sp. nov. (**Figure 4**, **Table 3**), *O. lunettae* sp. nov. exhibits a characteristic color gradient—from yellowish or cream at the base to more intensely colored tissue at the apex of the lobes—that mirrors the pattern typically observed in *O. lobularis* (**Figure 3 A to C’**). However, unlike *O. lobularis*, which is invariably blue, purple, or pink, *O. lunettae* sp. nov. consistently displays orange hues, making this gradient pattern and coloration combination diagnostic (**Figure 4G and G’**).

#### Morphological description

Like other *Oscarella* species, *O. lunettae* sp. nov. is an encrusting sponge characterized by irregular growth forms with broad, thick, elongated lobes (**Figure 4G and G’**, **Table 3**). It covers areas up to 80 cm^2^ with a thickness ranging from 10 to 20 mm. Numerous ostia, which are easily visible under low magnification, perforate the sponge surface giving to the sponge a rugous aspect (**Figure 4G and G’**), thus resembling *O. balibaloi* (**Figure 4B’ and C’**, **Table 3**), and *O viridis* (**Figure 4F’**, **Table 3**). The consistency of *O. lunettae* is relatively soft but not fragile, broadly comparable to that of *O. lobularis*.

#### Cytological description

The aquiferous system of this species appears similar to that described in other *Oscarella* species, with a sylleibid canal system, eurypylous choanocyte chambers, and a basal cavity. The ectosome thickness (4–40 µm) and the size of the largest choanocyte chambers (69–85 µm) fall within the ranges commonly observed in *Oscarella* species (**Supplementary Figure 4**, **Table 3**).

#### Ecology and distribution

Like other *Oscarella* species found in the Bay of Marseille, *O. lunettae* sp. nov. is a sciaphilous species, typically found in coralligenous environments. To date, it has been observed at only two locations: Levant and Tiboulen du Frioul. While *O. lobularis* and *O. tuberculata* are also present in these locations, *O. lunettae* has not been detected elsewhere, suggesting that it may occupy a slightly different ecological niche. All specimens were collected at depths between 30 and 40 m, suggesting that *O. lunettae* might thrive below the thermocline, where temperatures are more stable.

### *Oscarella sabinae* sp. nov. (Schenkelaars & Guiollot)

#### Holotype

*Muséum National d’Histoire Naturelle de Paris*, MNHN-IP-2019-3416, IMBE-B007-MAI95: Collected from France, Mediterranean Sea, Marseille, Maire Island (43.2096°N, 5.3353°W), at a depth of 8-18 m by Dorian Guillemain on 2024-06-19 (MAI_95, **Figure 3G and G’**). Specimen fixed in 4% paraformaldehyde and preserved in absolute methanol. No other material has been examined.

#### Molecular diagnosis

Molecular analysis of *O. sabinae* sp. nov. revealed distinct haplotypes for all mitochondrial markers, differentiating it from other species within the complex (**Supplementary Figure 2**). Comparative analysis of partial mitochondrial genomes identified 19 putative diagnostic positions unique to this species, including three in *cox1* (T_96_→C, T_336_→C and A_936_→G) and two in *nad1–atp8* (A_193_→G and A_251_→G). The quality of the ONT-derived partial mitochondrial genome was assessed based on sequencing coverage and consensus Phred quality, with high coverage and consistently high consensus quality across the assembly (**Supplementary Table 3**). The 19 diagnostic mitochondrial substitutions identified in *O. sabinae* were further checked against the underlying read data and showed high coverage and strong consensus support at each position (**Supplementary Table 4**), supporting their reliability as putative diagnostic characters. Two diagnostic substitutions were also detected in the *epic12* haplotype o when compared to all other haplotypes from species within the complex (G_14_→A and T_50_→A).

#### Morphological diagnosis

In the single specimen of *O. sabinae* sp. nov. examined to date, the lobes display an irregular surface that give the sponge a bright pink, cauliflower-like appearance (**Figure 3G and G’**). Although *O. veronicae* sp. nov. can also exhibit a cauliflower-like morphology (**Figure 3H and H’**, **Table 3**), none of the specimens observed showed a similarly intense pink coloration. Indeed, the hue of *O. sabinae* sp. nov. is more reminiscent of the pink form of *O. lobularis* (**Figure 3A and A’**), yet the latter species never exhibits a cauliflower-like surface (n = 42 in this study, with many more examined in the laboratory).

#### Morphological description

Like other *Oscarella* species, *O. sabinae* sp. nov. is an encrusting sponge characterized by irregular growth forms with thick, rounded lobes (**Table 3**). The specimen collected measured 10-15 cm^2^. While the specimen was extremely soft, it was not fragile.

#### Cytological description

The aquiferous system of this species appears similar to that described in other *Oscarella* species, with a sylleibid canal system, eurypylous choanocyte chambers, and a basal cavity (**Supplementary Figure 4**). The ectosome thickness (6–61 µm) and the size of the largest choanocyte chambers (56–68 µm) fall within the ranges commonly observed in *Oscarella* species (**Table 3**).

#### Ecology and Distribution

Out of 192 samples collected, only one specimen was identified as *O. sabinae* sp. nov., indicating that this species may be rare or that the sampling area may not represent its primary habitat. Consequently, it is premature to detail the ecology and distribution of this species based on the available data.

#### Remark

The specimen identified as *Oscarella rubra* by Gazave et al. (2013) may belong to the same species as the specimens examined here, given the identity of their *atp6* sequences. However, the name *O. rubra* is not applicable to an *Oscarella*, as it is currently treated as a synonym of *Aplysilla rubra* (Hanitsch, 1890).

### *Oscarella veronicae* sp. nov. (Schenkelaars & Guiollot)

#### Holotype

*Muséum National d’Histoire Naturelle de Paris*, MNHN-IP-2019-3417, IMBE-A088-MAI60: Collected from France, Mediterranean Sea, Marseille, Maire Island (43.2096°N, 5.3353°W), at a depth of 6-10 m by Dorian Guillemain on 2023-04-12 (MAI_35, **Figure 3H and H’**). Specimen fixed in 4% paraformaldehyde and preserved in absolute methanol.

#### Paratype

*Muséum National d’Histoire Naturelle de Paris*, MNHN-IP-2019-3418, IMBE-A081-MAI56: Collected from France, Mediterranean Sea, Marseille, Maire Island (43.2096°N, 5.3353°W), at a depth of 17-20 m by Dorian Guillemain on 2023-05-04 (MAI_56, **Supplementary Figure 1**). Specimen fixed in 4% paraformaldehyde and preserved in absolute methanol.

#### Other material examined

Station Marine d’Endoume, IMBE: 23 additional specimens collected from various locations across Marseille Bay, including 3PP (43.09°N, 05.36°W), Maïre Island (43.2096°N, 5.3353°W), Mugel (43.1001°N, 5.3618°W), Tiboulen du Frioul (43.2799°N, 5.2860°W), and Vesse (43.2026°N, 5.1539°W) (**Supplementary Table 1** and **Supplementary Figure 1**). Specimen fixed in 4% paraformaldehyde and preserved in absolute methanol.

#### Molecular diagnosis

Mitochondrial markers were found to differentiate *Oscarella veronicae* sp. nov. from related species, though they did not exhibit mutations exclusive to this species. However, comparison of partial mitochondrial genomes between *O. lobularis*, *O. tuberculata*, *O. veronicae* sp. nov., and *O. sabinae* sp. nov. identified a unique mutation specific to *O. veronicae* sp. nov. (A_7563_→G). This unique diagnostic mitochondrial substitution was further checked against the underlying read data and showed high coverage and strong consensus support (**Supplementary Table 4**). Furthermore, it was validated through PCR amplification across various specimens from each species (**Supplementary Figure 5**). In contrast, nuclear data provided a clearer distinction, with *epic12* haplotypes f and e both revealing four *O. veronicae* sp. nov. specific mutations despite the short length of the marker (T_29_→C, G_33_→A, A_38_→G, and T48→C) (**Supplementary Figure 2**).

#### Morphological diagnosis

Many specimens exhibited a cauliflower-like morphology closely resembling that of *O. sabinae* sp. nov. (**Figure 3G to H’**); however, unlike the latter, they never display a bright pink coloration (**Table 3**). Instead, most specimens exhibit a blue hue that differs from other species within the complex, appearing relatively species-specific: shades tend toward cyan, whereas *O. lobularis* and *O. tuberculata* display blues that are more grayish or purplish (**Figure 3C and F**).

#### Morphological description

Like other *Oscarella* species, *O. veronicae* sp. nov. is an encrusting sponge characterized by irregular growth forms with thick, rounded lobes (**Table 3**). It covers areas up to 20-30 cm^2^ and reaches a thickness of 10-15 mm. Most specimens display a bright or light blue coloration (**Figure 3D**), although exceptions include yellow (2 out of 25 specimens) or dark pink like *O. tuberculata* (one specimen, TIB_09, **Supplementary Figure 1, Table 3**). The sponge is neither soft nor cartilaginous in consistency.

#### Cytological description

The aquiferous system of this species appears similar to that described in other *Oscarella* species, with a sylleibid canal system, eurypylous choanocyte chambers, and a basal cavity (**Supplementary Figure 4**). The ectosome thickness (14–58 µm) falls within the ranges commonly observed in *Oscarella* species and tends to be rich in vacuolar cells (**Supplementary Figure 4D**, **Table 3**). Although the size of the largest choanocyte chambers (73–99 µm) also falls within the ranges commonly observed in *Oscarella* species, it is toward the upper end of the range (**Table 3**).

#### Ecology and distribution

*O. veronicae* sp. nov. is a sciaphilous species consistently found in sympatry with *O. tuberculata* and *O. lobularis*. It inhabits the cavities of coralligenous walls or the entrances of submarine caves at depths ranging from 5 to 35 m. To date, no data are available to confirm the presence or absence of this species elsewhere in the Mediterranean Sea or along the French Atlantic coasts.

## CONCLUSION

This study highlights the importance of integrating complementary genetic data and standardized methodologies in the identification and taxonomy of *Oscarella* species. By combining mitochondrial and nuclear markers with comprehensive photographic documentation, we refined species delimitation within this skeleton-less genus. Our results revealed four new species in the Bay of Marseille, underscoring both the hidden diversity of *Oscarella* and the evolutionary significance of color polymorphism. At the same time, we exposed major limitations in current practices, including the low resolution of the cytochrome c oxidase subunit I (*cox1*/*COI*) gene and the lack of genetic data for many accepted species. Addressing these gaps will require coordinated efforts to generate robust genetic frameworks, which are essential not only for reliable species identification but also for clarifying biogeographic patterns and evolutionary processes. Ultimately, advancing a standardized, data-rich taxonomy of *Oscarella* will strengthen our ability to assess, conserve, and understand marine biodiversity.

## ADDITIONAL INFORMATION

Authors declare no competing financial or non-financial interests in relation to the work described.

## ACKNOWLEDGEMENTS

Data was generated through the Molecular and Cell Biology Facility of IMBE, Aix-Marseille Univ, Avignon Univ, CNRS, IRD, Marseille. We are grateful to S. Chenesseau from the Morphology Facility of IMBE and to G. Deniel for preparing the semi-thin sections, as well as to the OSU Pytheas diving facility for their support in *Oscarella* sampling. We also warmly thank A. Chenuil-Maurel and A. Baumel for helpful input on manuscript preparation.

## DATA AVAILABILITY

The genetic datasets generated and analyzed during the current study are available in the NCBI repository, https://www.ncbi.nlm.nih.gov/. All accession numbers are provided in **Supplementary Table 2**. All other data generated during this study (pictures of animals) are included in this published article (and its Supplementary Information files).

## AUTHOR CONTRIBUTIONS

DG conducted the majority of the sampling. QS captured the images. EG and QS prepared the samples and performed all gene amplifications. QS carried out phylogenetic analysis and constructed haplotype networks. QS, ER, CB, and EG contributed to writing the paper. QS supervised the entire project.

