## Supplementary_Information for "Genetic and Morphological Insights Reveal the Hidden and Colorful Diversity in *Oscarella* Sponges"

**Supplementary Table 1: List of *Oscarella spp*. specimens.** It contains information regarding the sampling, the PCR results (haplotypes), and its assignment to a given species. (page 2)

**Supplementary Table 2: Accession numbers of mitochondrial genomes and haplotypes.** Species names with an asterisk correspond to partial mitochondrial genomes. Species names underlined in blue indicate those obtained in the present study. (page 5)

**Supplementary Table 3. Nanopore sequencing coverage and consensus quality statistics for mitochondrial amplicon assemblies.** Variants were detected using a minor allele frequency (MAF) threshold of ≥30% and a minimum read support of >30×, as defined by the Eurofins amplicon analysis pipeline. Phred quality scores represent the confidence assigned to the consensus base at each position, with higher scores indicating higher base-calling confidence. (page 6)

**Supplementary Table 4. Per-position Nanopore read support for phylogenetically relevant substitutions in mitochondrial assemblies.** (page 7)

**Supplementary Figure 1: Picture of each specimen.** Abbreviatons: 3PP, 3PP; JAR, Jarre; LEV, Levant; MAI, Maïre; MEJ, Mejean; MUG, Mugel; TIB, Tiboulen du Frioul; and VES, Vesse. (page 8)

**Supplementary Figure 2: Haplotype networks supporting the delimitation of four new *Oscarella* species.** Haplotype networks based on the *atp6*, *nad1-atp8*, *tatC*, *cox1* mitochondrial markers and *epic12* nuclear marker. Each haplotype (a, b, c, etc.) is depicted as a circle, with the size of the circle proportional to the number of specimens represented. Numbers between haplotypes indicate the number of mutations separating them. Colors correspond to species delimitation. Note that the *epic12* sequences were not phased. Yet, allelic variation was very limited in most species, with one to two alleles detected per species (except *O. tuberculata*), and a small number of heterozygous individuals were identified in which both allelic variants could be unambiguously recognized. (page 16)

**Supplementary Figure 3: Maximum likelihood (ML) phylogenetic tree of *Oscarella* species based on eight mitochondrial protein-coding genes (*cox1*, *nad1*, *atp8*, *atp6*, *cox3*, *atp9*, *nad4*, and *nad6*).** Only nodes with bootstrap values greater than 85% (1000 BS replicates) are shown. Dots on the tree indicate the inferred ancestral color state of the common ancestor of *Oscarella* and the last common ancestor of *O. lobularis*, *O. sabinae* sp. nov., *O. tuberculata*, and *O. veronicae* sp. nov. Note that *Pseudocorticium jarrei* is proposed to be renamed *Oscarella jarrei*. (page 17)

**Supplementary Figure 4. Histological semi-thin sections of the four *Oscarella* species newly described in this study.** (A) *Oscarella fragilis* sp. nov.; (B) *Oscarella lunettae* sp. nov.; (C) *Oscarella sabinae* sp. nov.; (D) *Oscarella veronicae* sp. nov. Abbreviations: c.c., choanocyte chamber; ex, exopinacocyte; i.c., inhalant canal; v.c., vacuolar cell. (page 18)

**Supplementary Figure 5: Sequence alignment highlighting the unique diagnostic mutation found in the partial mitochondrial genome of *O. veronicae* sp. nov.** This sequence, we called *SNP*, was amplified and sequenced for several specimens belonging to the species complex. The mutation specific to *O. veronicae* sp. nov. is marked by a black arrow. (page 19)

Sup. Table 1

| | Specimen | | | | | | PCRs (haplotypes) | | | | | Conclusion |
|---|---|---|---|---|---|---|---|---|---|---|---|---|
| | Name | Year | Month | Day | Where | Depth | *atp6* | *nad1-atp8* | *epic12* | *tatC* | *cox1* | Species |
| 1 | MAI_42 | 2023 | 4 | 12 | Maïre | 6-10m | a | a | a | b | | tuberculata |
| 2 | MAI_43 | 2023 | 4 | 12 | Maïre | 6-10m | a | a | a | b | | tuberculata |
| 3 | MAI_65 | 2023 | 5 | 4 | Maïre | 17-20m | a | a | a | | b | tuberculata |
| 4 | MEJ_06 | 2023 | 5 | 4 | Mejean | 25-30m | a | a | a | | b | tuberculata |
| 5 | JAR_08 | 2023 | 4 | 26 | Jarre | 14-17m | a | a | a | | | tuberculata |
| 6 | MUG_09 | 2023 | 5 | 29 | Mugel | 3-10m | a | a | a | | | tuberculata |
| 7 | MUG_10 | 2023 | 5 | 29 | Mugel | 3-10m | a | a | a | | | tuberculata |
| 8 | VES_05 | 2023 | 5 | 24 | Vesse | 16m | a | a | a | | | tuberculata |
| 9 | VES_06 | 2023 | 5 | 24 | Vesse | 16m | a | a | a | | | tuberculata |
| 10 | VES_10 | 2023 | 5 | 24 | Vesse | 16m | a | a | a | | | tuberculata |
| 11 | VES_13 | 2023 | 5 | 24 | Vesse | 16m | a | a | a | | | tuberculata |
| 12 | VES_15 | 2023 | 5 | 24 | Vesse | 16m | a | a | a | | | tuberculata |
| 13 | MAI_07 | 2022 | 12 | 7 | Maïre | ??? | a | a | | b | | tuberculata |
| 14 | MAI_09 | 2022 | 12 | 7 | Maïre | ??? | a | a | | b | | tuberculata |
| 15 | MAI_10 | 2022 | 12 | 7 | Maïre | ??? | a | a | | b | | tuberculata |
| 16 | MAI_45 | 2023 | 4 | 12 | Maïre | 6-10m | a | a | | b | | tuberculata |
| 17 | MAI_66 | 2023 | 5 | 4 | Maïre | 17-20m | a | a | | b | | tuberculata |
| 18 | MAI_18 | 2023 | 3 | 9 | Maïre | ??? | a | | a | b | | tuberculata |
| 19 | MAI_21 | 2023 | 3 | 9 | Maïre | ??? | a | | a | b | | tuberculata |
| 20 | MAI_22 | 2023 | 3 | 9 | Maïre | ??? | a | | a | b | | tuberculata |
| 21 | MEJ_05 | 2023 | 5 | 4 | Mejean | 25-30m | a | a | | | | tuberculata |
| 22 | MUG_04 | 2023 | 5 | 29 | Mugel | 3-10m | a | a | | | | tuberculata |
| 23 | TIB_03 | 2023 | 6 | 28 | Tiboulen | 30-35m | a | a | | | | tuberculata |
| 24 | TIB_12 | 2023 | 6 | 28 | Tiboulen | 30-35m | a | a | | | | tuberculata |
| 25 | TIB_14 | 2023 | 6 | 28 | Tiboulen | 30-35m | a | a | | | | tuberculata |
| 26 | VES_01 | 2023 | 5 | 24 | Vesse | 16m | a | a | | | | tuberculata |
| 27 | VES_04 | 2023 | 5 | 24 | Vesse | 16m | a | a | | | | tuberculata |
| 28 | VES_07 | 2023 | 5 | 24 | Vesse | 16m | a | a | | | | tuberculata |
| 29 | VES_11 | 2023 | 5 | 24 | Vesse | 16m | a | a | | | | tuberculata |
| 30 | VES_09 | 2023 | 5 | 24 | Vesse | 16m | a | | a | | | tuberculata |
| 31 | MAI_08 | 2022 | 12 | 7 | Maïre | ??? | a | | | b | | tuberculata |
| 32 | MAI_69 | 2023 | 7 | 12 | Maïre | 8-18m | a | | | | | tuberculata |
| 33 | MAI_70 | 2023 | 7 | 12 | Maïre | 8-18m | a | | | | | tuberculata |
| 34 | MAI_71 | 2023 | 7 | 12 | Maïre | 8-18m | a | | | | | tuberculata |
| 35 | MAI_72 | 2023 | 7 | 12 | Maïre | 8-18m | a | | | | | tuberculata |
| 36 | MAI_73 | 2023 | 7 | 12 | Maïre | 8-18m | a | | | | | tuberculata |
| 37 | MAI_74 | 2023 | 7 | 12 | Maïre | 8-18m | a | | | | | tuberculata |
| 38 | MAI_76 | 2023 | 7 | 12 | Maïre | 8-18m | a | | | | | tuberculata |
| 39 | MAI_79 | 2023 | 7 | 12 | Maïre | 8-18m | a | | | | | tuberculata |
| 40 | MAI_81 | 2023 | 7 | 12 | Maïre | 8-18m | a | | | | | tuberculata |
| 41 | MAI_82 | 2023 | 7 | 12 | Maïre | 8-18m | a | | | | | tuberculata |
| 42 | MAI_85 | 2023 | 7 | 12 | Maïre | 8-18m | a | | | | | tuberculata |
| 43 | MAI_86 | 2023 | 7 | 12 | Maïre | 8-18m | a | | | | | tuberculata |
| 44 | MAI_87 | 2023 | 7 | 12 | Maïre | 8-18m | a | | | | | tuberculata |
| 45 | MAI_94 | 2023 | 7 | 12 | Maïre | 8-18m | a | | | | | tuberculata |
| 46 | MEJ_08 | 2023 | 5 | 4 | Mejean | 25-30m | a | | | | | tuberculata |
| 47 | MEJ_09 | 2023 | 5 | 4 | Mejean | 25-30m | a | | | | | tuberculata |
| 48 | TIB_05 | 2023 | 6 | 28 | Tiboulen | 30-35m | a | | | | | tuberculata |
| 49 | TIB_15 | 2023 | 6 | 28 | Tiboulen | 30-35m | a | | | | | tuberculata |
| 50 | TIB_16 | 2023 | 6 | 28 | Tiboulen | 30-35m | a | | | | | tuberculata |
| 51 | VES_02 | 2023 | 5 | 24 | Vesse | 16m | a | | | | | tuberculata |
| 52 | VES_03 | 2023 | 5 | 24 | Vesse | 16m | a | | | | | tuberculata |
| 53 | VES_12 | 2023 | 5 | 24 | Vesse | 16m | a | | | | | tuberculata |
| 54 | VES_22 | 2023 | 7 | 10 | Vesse | 13-15m | a | | | | | tuberculata |
| 55 | VES_23 | 2023 | 7 | 10 | Vesse | 13-15m | a | | | | | tuberculata |
| 56 | VES_24 | 2023 | 7 | 10 | Vesse | 13-15m | a | | | | | tuberculata |
| 57 | VES_25 | 2023 | 7 | 10 | Vesse | 13-15m | a | | | | | tuberculata |
| 58 | VES_26 | 2023 | 7 | 10 | Vesse | 13-15m | a | | | | | tuberculata |
| 59 | VES_28 | 2023 | 7 | 10 | Vesse | 13-15m | a | | | | | tuberculata |
| 60 | VES_29 | 2023 | 7 | 10 | Vesse | 13-15m | a | | | | | tuberculata |
| 61 | VES_30 | 2023 | 7 | 10 | Vesse | 13-15m | a | | | | | tuberculata |
| 62 | VES_32 | 2023 | 7 | 10 | Vesse | 13-15m | a | | | | | tuberculata |
| 63 | MAI_32 | 2023 | 4 | 12 | Maïre | 6-10m | a | a | c | b | b | tuberculata |
| 64 | MUG_02 | 2023 | 5 | 29 | Mugel | 3-10m | a | a | c | | b | tuberculata |

| | | | | | | | | | | | | |
|---|---|---|---|---|---|---|---|---|---|---|---|---|
| 65 | TIB_02 | 2023 | 6 | 28 | Tiboulen | 30-35m | a | a | c | | b | tuberculata |
| 66 | VES_19 | 2023 | 5 | 24 | Vesse | 16m | a | a | c | | | tuberculata |
| 67 | MAI_44 | 2023 | 4 | 12 | Maïre | 6-10m | a | a | h | b | | tuberculata |
| 68 | VES_08 | 2023 | 5 | 24 | Vesse | 16m | a | a | l | | b | tuberculata |
| 69 | MEJ_07 | 2023 | 5 | 4 | Mejean | 25-30m | a | a | m | | b | tuberculata |
| 70 | MAI_41 | 2023 | 4 | 12 | Maïre | 6-10m | a | | k | b | | tuberculata |
| 71 | JAR_16 | 2023 | 4 | 26 | Jarre | 14-17m | a | h | c | | | tuberculata |
| 72 | VES_27 | 2023 | 7 | 10 | Vesse | 13-15m | a | h | | | | tuberculata |
| 73 | JAR_04 | 2023 | 4 | 26 | Jarre | 14-17m | d | b | c | | d | tuberculata |
| 74 | JAR_05 | 2023 | 4 | 26 | Jarre | 14-17m | d | b | c | | | tuberculata |
| 75 | JAR_12 | 2023 | 4 | 26 | Jarre | 14-17m | d | b | c | | | tuberculata |
| 76 | MAI_64 | 2023 | 5 | 4 | Maïre | 17-20m | d | b | | | | tuberculata |
| 77 | JAR_09 | 2023 | 4 | 26 | Jarre | 14-17m | d | | | | | tuberculata |
| 78 | MAI_75 | 2023 | 7 | 12 | Maïre | 8-18m | d | | | | | tuberculata |
| 79 | MAI_77 | 2023 | 7 | 12 | Maïre | 8-18m | d | | | | | tuberculata |
| 80 | MAI_78 | 2023 | 7 | 12 | Maïre | 8-18m | d | | | | | tuberculata |
| 81 | MAI_80 | 2023 | 7 | 12 | Maïre | 8-18m | d | | | | | tuberculata |
| 82 | MAI_83 | 2023 | 7 | 12 | Maïre | 8-18m | d | | | | | tuberculata |
| 83 | MAI_84 | 2023 | 7 | 12 | Maïre | 8-18m | d | | | | | tuberculata |
| 84 | MAI_11 | 2022 | 12 | 7 | Maïre | ??? | d | b | a | d | | tuberculata |
| 85 | MAI_23 | 2023 | 3 | 9 | Maïre | ??? | d | b | a | d | | tuberculata |
| 86 | MEJ_10 | 2023 | 5 | 4 | Mejean | 25-30m | d | b | a | d | | tuberculata |
| 87 | JAR_02 | 2023 | 4 | 26 | Jarre | 14-17m | d | b | a | | | tuberculata |
| 88 | JAR_10 | 2023 | 4 | 26 | Jarre | 14-17m | d | b | a | | | tuberculata |
| 89 | MAI_63 | 2023 | 5 | 4 | Maïre | 17-20m | d | b | a | | | tuberculata |
| 90 | JAR_03 | 2023 | 4 | 26 | Jarre | 14-17m | d | | a | | | tuberculata |
| 91 | MAI_37 | 2023 | 4 | 12 | Maïre | 6-10m | d | | h | d | | tuberculata |
| 92 | JAR_06 | 2023 | 4 | 26 | Jarre | 14-17m | d | b | n | d | d | tuberculata |
| 93 | MAI_67 | 2023 | 5 | 4 | Maïre | 17-20m | d | b | a | b | | tuberculata |
| 94 | MAI_38 | 2023 | 4 | 12 | Maïre | 6-10m | e | a | c | b | | tuberculata |
| 95 | MAI_39 | 2023 | 4 | 12 | Maïre | 6-10m | e | a | c | b | | tuberculata |
| 96 | MAI_40 | 2023 | 4 | 12 | Maïre | 6-10m | e | a | c | b | | tuberculata |
| 97 | MUG_01 | 2023 | 5 | 29 | Mugel | 3-10m | e | b | a | | | tuberculata |
| 98 | MUG_06 | 2023 | 5 | 29 | Mugel | 3-10m | e | b | a | | | tuberculata |
| 99 | VES_18 | 2023 | 5 | 24 | Vesse | 16m | e | b | a | | | tuberculata |
| 100 | VES_21 | 2023 | 7 | 10 | Vesse | 13-15m | e | b | a | | | tuberculata |
| 101 | MEJ_04 | 2023 | 5 | 4 | Mejean | 25-30m | e | b | a | f | | tuberculata |
| 102 | MUG_07 | 2023 | 5 | 29 | Mugel | 3-10m | e | b | c | | | tuberculata |
| 103 | MUG_08 | 2023 | 5 | 29 | Mugel | 3-10m | e | b | c | | | tuberculata |
| 104 | VES_14 | 2023 | 5 | 24 | Vesse | 16m | j | a | a | | | tuberculata |
| 105 | MAI_95 | 2024 | 6 | 19 | Maïre | 8-18m | k | p | o | g | e | sabinae |
| 106 | VES_16 | 2023 | 5 | 24 | Vesse | 16m | c | e | e | c | c | veronicae |
| 107 | MAI_61 | 2023 | 5 | 4 | Maïre | 17-20m | c | e | | c | | veronicae |
| 108 | MAI_34 | 2023 | 4 | 12 | Maïre | 6-10m | c | | e | c | | veronicae |
| 109 | MAI_12 | 2022 | 12 | 7 | Maïre | ??? | c | | | c | | veronicae |
| 110 | MAI_19 | 2023 | 3 | 9 | Maïre | ??? | c | | | c | | veronicae |
| 111 | MAI_36 | 2023 | 4 | 12 | Maïre | 6-10m | c | | | c | | veronicae |
| 112 | MAI_56 | 2023 | 5 | 4 | Maïre | 17-20m | c | | | c | | veronicae |
| 113 | MAI_62 | 2023 | 5 | 4 | Maïre | 17-20m | c | | | c | | veronicae |
| 114 | MAI_68 | 2023 | 5 | 4 | Maïre | 17-20m | c | e | | | | veronicae |
| 115 | VES_31 | 2023 | 7 | 10 | Vesse | 13-15m | c | e | | | | veronicae |
| 116 | MAI_88 | 2023 | 7 | 12 | Maïre | 8-18m | c | | | | | veronicae |
| 117 | MAI_89 | 2023 | 7 | 12 | Maïre | 8-18m | c | | | | | veronicae |
| 118 | MAI_90 | 2023 | 7 | 12 | Maïre | 8-18m | c | | | | | veronicae |
| 119 | MAI_91 | 2023 | 7 | 12 | Maïre | 8-18m | c | | | | | veronicae |
| 120 | MAI_92 | 2023 | 7 | 12 | Maïre | 8-18m | c | | | | | veronicae |
| 121 | MAI_93 | 2023 | 7 | 12 | Maïre | 8-18m | c | | | | | veronicae |
| 122 | MAI_57 | 2023 | 5 | 4 | Maïre | 17-20m | c | e | e | c | a | veronicae |
| 123 | VES_17 | 2023 | 5 | 24 | Vesse | 16m | c | e | e/f | c | a | veronicae |
| 124 | MAI_05 | 2022 | 12 | 7 | Maïre | ??? | c | | e | c | a | veronicae |
| 125 | MAI_60 | 2023 | 5 | 4 | Maïre | 17-20m | c | e | f | c | | veronicae |
| 126 | MUG_03 | 2023 | 5 | 29 | Mugel | 3-10m | c | e | f | c | | veronicae |
| 127 | 3PP_01 | 2024 | 1 | 31 | 3PP | 17m | c | | f | | c | veronicae |
| 128 | MAI_33 | 2023 | 4 | 12 | Maïre | 6-10m | c | | f | c | a | veronicae |
| 129 | MAI_35 | 2023 | 4 | 12 | Maïre | 6-10m | c | | f | c | a | veronicae |
| 130 | TIB_09 | 2023 | 6 | 28 | Tiboulen | 30-35m | c | m | e | c | a | veronicae |
| 131 | MAI_01 | 2022 | 9 | 5 | Maïre | ??? | b | c | b | a | a | lobularis |
| 132 | MUG_05 | 2023 | 5 | 29 | Mugel | 3-10m | b | c | b | a | a | lobularis |
| 133 | VES_20 | 2023 | 7 | 10 | Vesse | 13-15m | b | c | b | a | a | lobularis |

| | | | | | | | | | | | | |
|---|---|---|---|---|---|---|---|---|---|---|---|---|
| 134 | MAI_15 | 2023 | 3 | 9 | Maïre | ??? | b | c | b | a | | lobularis |
| 135 | MAI_17 | 2023 | 3 | 9 | Maïre | ??? | b | c | b | a | | lobularis |
| 136 | MAI_20 | 2023 | 3 | 9 | Maïre | ??? | b | c | b | a | | lobularis |
| 137 | MAI_25 | 2023 | 4 | 12 | Maïre | 6-10m | b | c | b | a | | lobularis |
| 138 | MAI_26 | 2023 | 4 | 12 | Maïre | 6-10m | b | c | b | a | | lobularis |
| 139 | MAI_27 | 2023 | 4 | 12 | Maïre | 6-10m | b | c | b | a | | lobularis |
| 140 | MAI_28 | 2023 | 4 | 12 | Maïre | 6-10m | b | c | b | a | | lobularis |
| 141 | MEJ_02 | 2023 | 5 | 4 | Mejean | 25-30m | b | c | b | a | | lobularis |
| 142 | MAI_06 | 2022 | 12 | 7 | Maïre | ??? | b | c | | a | a | lobularis |
| 143 | MAI_13 | 2023 | 3 | 9 | Maïre | ??? | b | c | | a | | lobularis |
| 144 | MAI_14 | 2023 | 3 | 9 | Maïre | ??? | b | c | | a | | lobularis |
| 145 | MAI_16 | 2023 | 3 | 9 | Maïre | ??? | b | c | | a | | lobularis |
| 146 | MAI_55 | 2023 | 5 | 4 | Maïre | 17-20m | b | c | | a | | lobularis |
| 147 | MAI_58 | 2023 | 5 | 4 | Maïre | 17-20m | b | c | | | a | lobularis |
| 148 | MAI_52 | 2023 | 5 | 4 | Maïre | 17-20m | b | c | | | | lobularis |
| 149 | MAI_59 | 2023 | 5 | 4 | Maïre | 17-20m | b | c | | | | lobularis |
| 150 | MAI_31 | 2023 | 4 | 12 | Maïre | 6-10m | b | | | a | | lobularis |
| 151 | TIB_17 | 2023 | 6 | 28 | Tiboulen | 30-35m | b | | | | | lobularis |
| 152 | MAI_02 | 2022 | 9 | 5 | Maïre | ??? | b | d | b | a | | lobularis |
| 153 | MAI_03 | 2022 | 9 | 5 | Maïre | ??? | b | d | b | a | | lobularis |
| 154 | MAI_24 | 2023 | 4 | 12 | Maïre | 6-10m | b | d | b | a | | lobularis |
| 155 | MAI_29 | 2023 | 4 | 12 | Maïre | 6-10m | b | d | b | a | | lobularis |
| 156 | MAI_53 | 2023 | 5 | 4 | Maïre | 17-20m | b | d | b | a | | lobularis |
| 157 | MAI_54 | 2023 | 5 | 4 | Maïre | 17-20m | b | d | b | a | | lobularis |
| 158 | TIB_04 | 2023 | 6 | 28 | Tiboulen | 30-35m | b | d | b | | a | lobularis |
| 159 | TIB_07 | 2023 | 6 | 28 | Tiboulen | 30-35m | b | d | b | | a | lobularis |
| 160 | TIB_13 | 2023 | 6 | 28 | Tiboulen | 30-35m | b | d | b | | a | lobularis |
| 161 | MAI_04 | 2022 | 12 | 7 | Maïre | ??? | b | d | | a | | lobularis |
| 162 | MAI_30 | 2023 | 4 | 12 | Maïre | 6-10m | b | d | | a | | lobularis |
| 163 | MAI_46 | 2023 | 5 | 4 | Maïre | 17-20m | b | d | | a | | lobularis |
| 164 | MAI_47 | 2023 | 5 | 4 | Maïre | 17-20m | b | d | | a | | lobularis |
| 165 | MAI_48 | 2023 | 5 | 4 | Maïre | 17-20m | b | d | | a | | lobularis |
| 166 | MAI_49 | 2023 | 5 | 4 | Maïre | 17-20m | b | d | | a | | lobularis |
| 167 | MAI_50 | 2023 | 5 | 4 | Maïre | 17-20m | b | d | | a | | lobularis |
| 168 | MAI_51 | 2023 | 5 | 4 | Maïre | 17-20m | b | d | | a | | lobularis |
| 169 | TIB_06 | 2023 | 6 | 28 | Tiboulen | 30-35m | b | d | | | | lobularis |
| 170 | TIB_08 | 2023 | 6 | 28 | Tiboulen | 30-35m | b | d | | | | lobularis |
| 171 | MEJ_01 | 2023 | 5 | 4 | Mejean | 25-30m | b | i | b | a | a | lobularis |
| 172 | MEJ_03 | 2023 | 5 | 4 | Mejean | 25-30m | b | i | b | a | | lobularis |
| 173 | 3PP_04 | 2024 | 1 | 31 | 3PP | 17m | g | j | j | | | viridis like |
| 174 | JAR_13 | 2024 | 1 | 16 | Jarre | ??? | g | j | j | | | viridis like |
| 175 | LEV_04 | 2024 | 3 | 7 | Levant | 40m | f | g | g | | | lunettae |
| 176 | LEV_05 | 2024 | 3 | 7 | Levant | 40m | f | g | g | | | lunettae |
| 177 | TIB_01 | 2023 | 6 | 28 | Tiboulen | 30-35m | f | g | g | | | lunettae |
| 178 | TIB_10 | 2023 | 6 | 28 | Tiboulen | 30-35m | f | g | g | | | lunettae |
| 179 | LEV_03 | 2024 | 3 | 7 | Levant | 40m | f | n | g | | | lunettae |
| 180 | TIB_11 | 2023 | 6 | 28 | Tiboulen | 30-35m | i | l | i | | | fragilis |
| 181 | JAR_01 | 2023 | 4 | 26 | Jarre | 14-17m | i | l | i / p | | | fragilis |
| 182 | JAR_15 | 2023 | 4 | 26 | Jarre | 14-17m | no | f | d | e | | balibaloi |
| 183 | MEJ_11 | 2023 | 5 | 4 | Mejean | 25-30m | no | f | d | e | | balibaloi |
| 184 | MEJ_12 | 2023 | 5 | 4 | Mejean | 25-30m | no | f | d | e | | balibaloi |
| 185 | 3PP_02 | 2024 | 1 | 31 | 3PP | 17m | no | f | d | | | balibaloi |
| 186 | 3PP_03 | 2024 | 1 | 31 | 3PP | 17m | no | f | d | | | balibaloi |
| 187 | 3PP_05 | 2024 | 1 | 31 | 3PP | 17m | no | f | d | | | balibaloi |
| 188 | LEV_01 | 2024 | 3 | 7 | Levant | 40m | no | | d | | | balibaloi |
| 189 | LEV_02 | 2024 | 3 | 7 | Levant | 40m | no | | d | | | balibaloi |
| 190 | JAR_07 | 2023 | 4 | 26 | Jarre | 14-17m | no | o | d | e | | balibaloi |
| 191 | JAR_11 | 2023 | 4 | 26 | Jarre | 14-17m | h | k | no | | | microlobata |
| 192 | JAR_14 | 2024 | 1 | 16 | Jarre | ??? | h | k | no | | | microlobata |

Sup. Table 2

| | Accession (NCBI) | | | | | |
|---|---|---|---|---|---|---|
| Scientific Name | mtGenome | *atp6* | *cox1* | *nad1-atp8* | *tatC* | *epic12* |
| *Corticium candelabrum* | HQ269363.1 | n.a. | n.a. | n.a. | n.a. | n.a. |
| *Oscarella balibaloi** | KY682865.1 | PCR failed | n.a. | PX242164 and PX242173 | PX252138 | PX262468 |
| *Oscarella carmela* | EF081250.1 | n.a. | n.a. | n.a. | n.a. | n.a. |
| *Oscarella fragilis sp. nov.** | PX112336 | PX242156 | n.a. | PX242170 | n.a. | PX262473 and PX262480 |
| *Oscarella lobularis* | OX382170.1 | PX242148 | PX261001 | PX242161, PX242162 and PX242167 | PX252134 | PX262466 |
| *Oscarella lunettae sp. nov.** | PX112335 | PX242153 | n.a. | PX242165 and PX242172 | n.a. | PX262471 |
| *Oscarella malakhovi* | HQ269364 | n.a. | n.a. | n.a. | n.a. | n.a. |
| *Oscarella microlobata* | HQ269355.1 | PX242155 | n.a. | PX242169 | n.a. | PCR failed |
| *Oscarella pearsei* | KY682864.1 | n.a. | n.a. | n.a. | n.a. | n.a. |
| *Oscarella sabinae sp. nov.** | PX112334 | PX242158 | PX261006 | PX242174 | PX252140 | PX262479 |
| *Oscarella tuberculata* | HQ269353.1 and JX963640.1 | PX242147, PX242150, PX242151, PX242152 and PX242157 | PX261003 and PX261005 | PX242159, PX242160 and PX242166 | PX252135, PX252137 and PX252139 | PX262465, PX262467, PX262472, PX262475, PX262476, PX262477, and PX262478 |
| *Oscarella veronicae sp. nov.** | PX112337 | PX242149 | PX261002 and PX261004 | PX242163 and PX242171 | PX252136 | PX262469 and PX262470 |
| *Oscarella viridis* | HQ269358.1 | PX242154 | n.a. | PX242168 | n.a. | PX262474 |
| *Plakortis simplex* | HQ269362.1 | n.a. | n.a. | n.a. | n.a. | n.a. |
| *Pseudocorticium jarrei* | HQ269357.1 | n.a. | n.a. | n.a. | n.a. | n.a. |

Sup. Table 3

| | | | | Coverage | | | | Phred | | | | |
|---|---|---|---|---|---|---|---|---|---|---|---|---|
| Species | Specimen | Length (bp) | Total reads | 5th percentil | Med | Mean | Max | Min | Med | Mean | Max | Variants detected |
| *Oscarella fragilis* | JAR_01 | 9,890 | 3,015× | 358× | 408× | 416× | 601× | 28 | 40 | 40 | 40 | No |
| Oscarella lunettae | LEV_05 | 9,967 | 2,842× | 68× | 91× | 150× | 482× | 29 | 40 | 40 | 40 | No |
| *Oscarella sabinae* | MAI_95 | 9,879 | 6,622× | 499× | 540× | 777× | 1,930× | 28 | 40 | 40 | 40 | No |
| *Oscarella veronicae* | MAI_33 | 9,949 | 3,793× | 425× | 555× | 560× | 2,628× | 29 | 40 | 40 | 40 | No |

Sup. Table 4

| Species | Specimen | NCBI position | Consensus base | Coverage | Consensus support | Major alternative | Alternative support |
|---|---|---|---|---|---|---|---|
| *Oscarella sabinae* | MAI_95 | 135 | G | 964× | 957× | A | 5× |
| *Oscarella sabinae* | MAI_95 | 725 | C | 1,326× | 1,316× | T | 3× |
| *Oscarella sabinae* | MAI_95 | 1,325 | G | 823× | 815× | A and T | 3× |
| *Oscarella sabinae* | MAI_95 | 1,565 | G | 652× | 644× | A and C | 2× |
| *Oscarella sabinae* | MAI_95 | 2,186 | G | 555× | 555× | n.a. | n.a. |
| *Oscarella sabinae* | MAI_95 | 2,605 | T | 547× | 547× | n.a. | n.a. |
| *Oscarella sabinae* | MAI_95 | 3,451 | C | 535× | 532× | T | 3× |
| *Oscarella sabinae* | MAI_95 | 3,509 | C | 534× | 533× | T | 1× |
| *Oscarella sabinae* | MAI_95 | 4,612 | A | 528× | 524× | C | 2× |
| *Oscarella sabinae* | MAI_95 | 5,228 | A | 522× | 515× | G | 6× |
| *Oscarella sabinae* | MAI_95 | 5,327 | G | 523× | 522× | A | 1× |
| *Oscarella sabinae* | MAI_95 | 5,343 | G | 521× | 517× | A | 2× |
| *Oscarella sabinae* | MAI_95 | 5,390 | G | 522× | 518× | T | 3× |
| *Oscarella sabinae* | MAI_95 | 5,837 | C | 496× | 492× | A and T | 1× |
| *Oscarella sabinae* | MAI_95 | 6,424 | A | 512× | 509× | C | 1× |
| *Oscarella sabinae* | MAI_95 | 6,699 | G | 498× | 497× | n.a. | n.a. |
| *Oscarella sabinae* | MAI_95 | 7,264 | G | 505× | 503× | A | 1× |
| *Oscarella sabinae* | MAI_95 | 8,521 | C | 1,286× | 1272× | A | 11× |
| *Oscarella sabinae* | MAI_95 | 9,761 | G | 935 | 928× | A | 4× |
| *Oscarella veronicae* | MAI_33 | 7,563 | G | 626 | 610× | C | 2× |

3PP_01
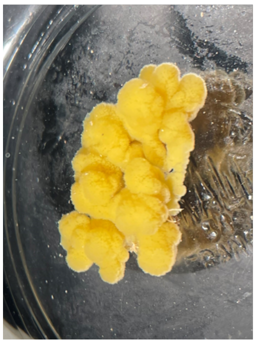
3PP_02
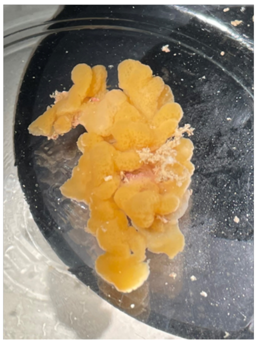
3PP_03
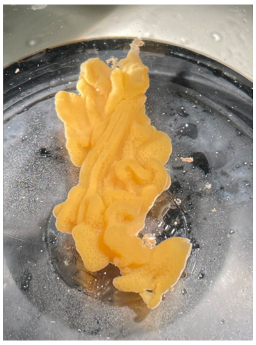
3PP_04
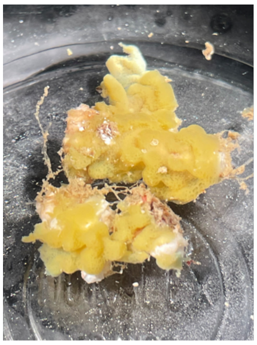
3PP_05
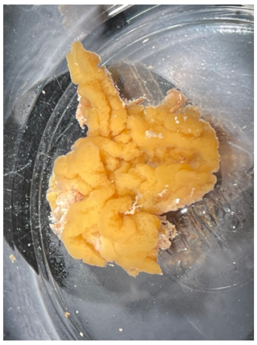
JAR_01
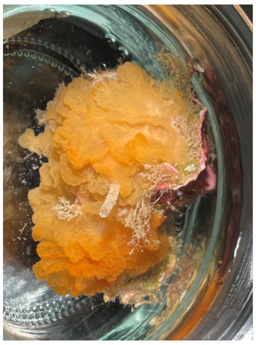
JAR_02
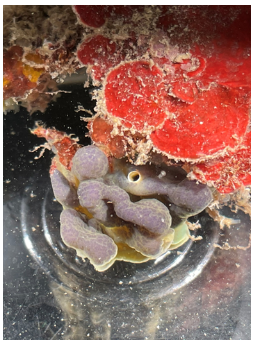
JAR_03
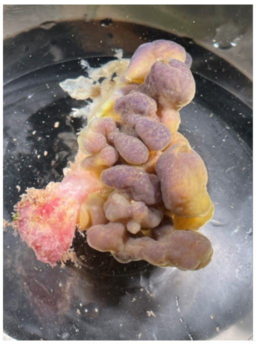
JAR_04
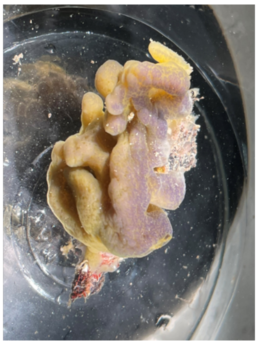
JAR_05
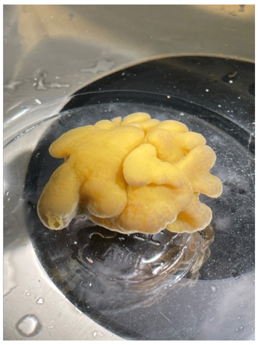
JAR_06
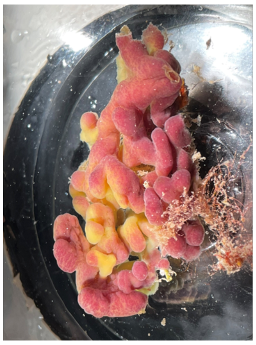
JAR_07
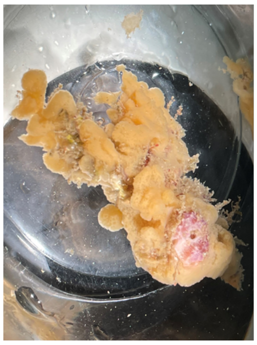
JAR_08
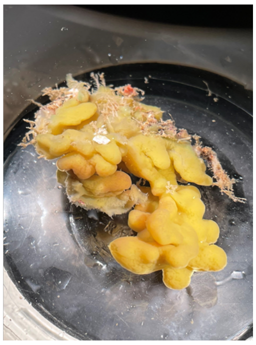
JAR_09
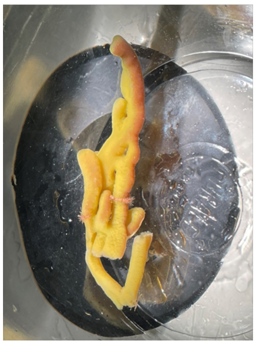
JAR_10
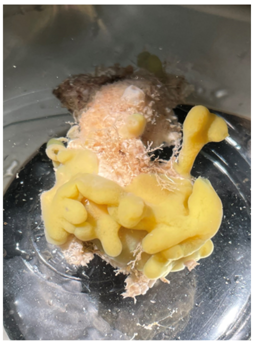
JAR_11
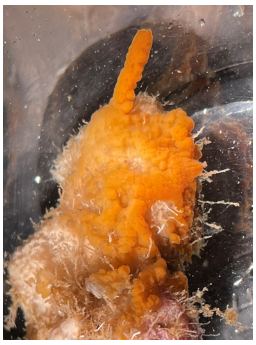
JAR_12
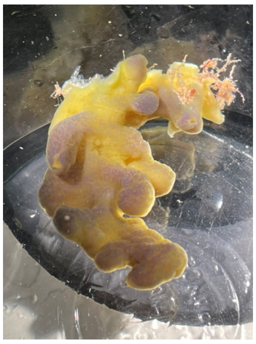
JAR_13
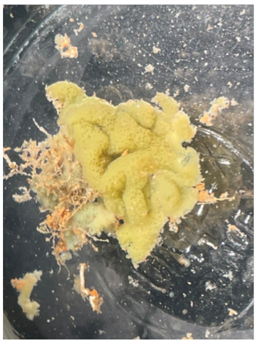
JAR_14
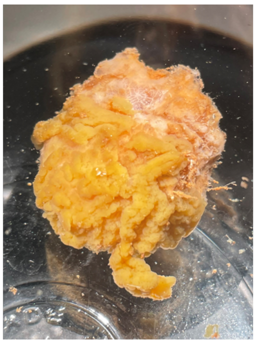
JAR_15
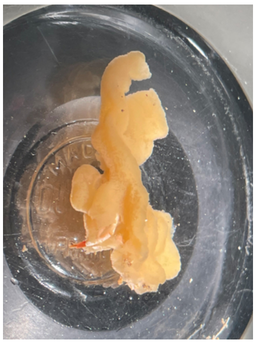
JAR_16
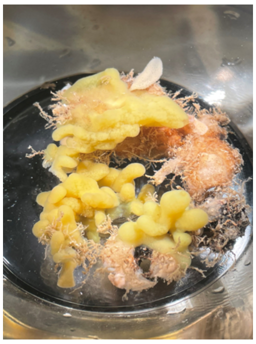
LEV_01
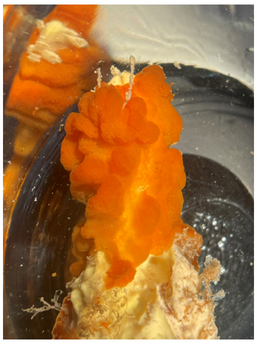
LEV_02
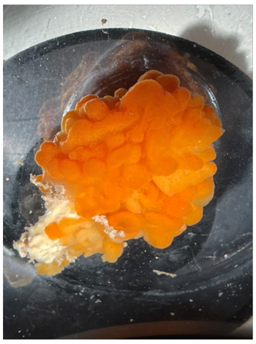
LEV_03
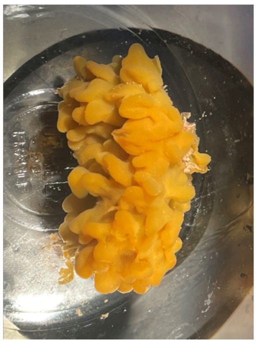
LEV_04
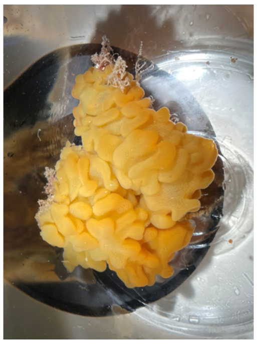

LEV_05

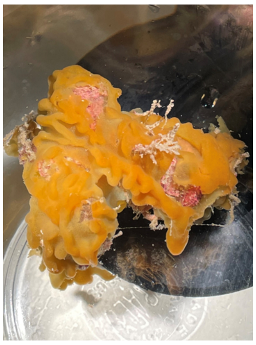

MAI_01

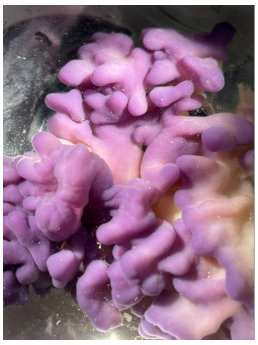

MAI_02

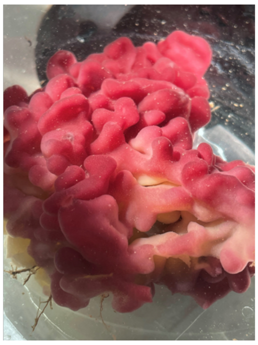

MAI_03

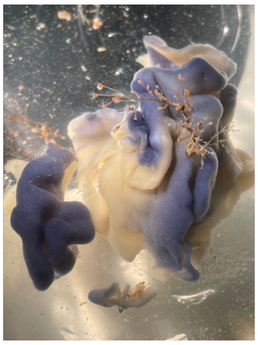

MAI_04

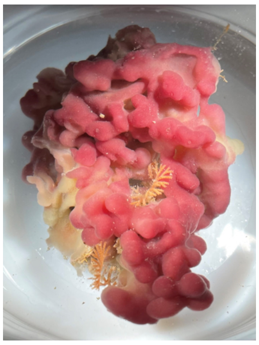

MAI_05

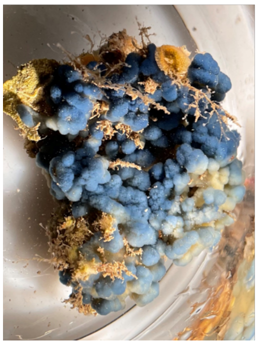

MAI_06

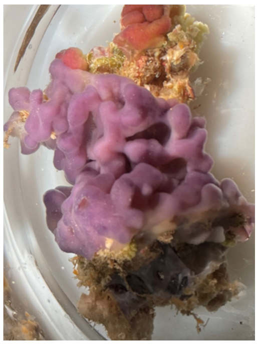

MAI_07

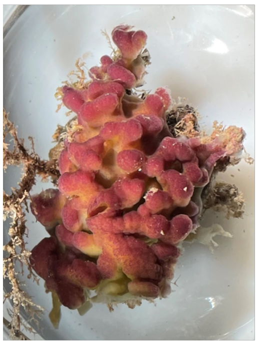

MAI_08

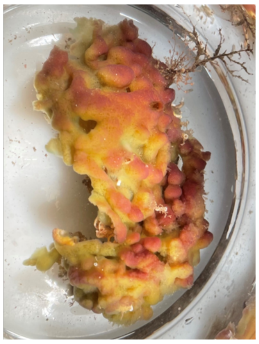

MAI_09

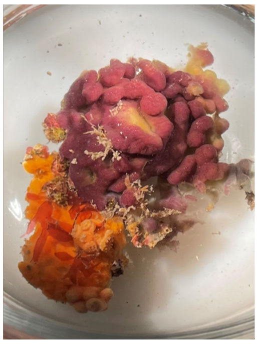

MAI_10

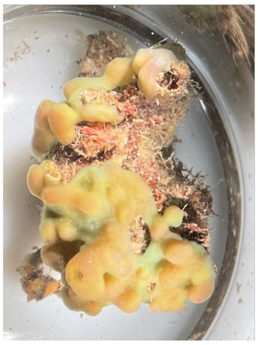

MAI_11

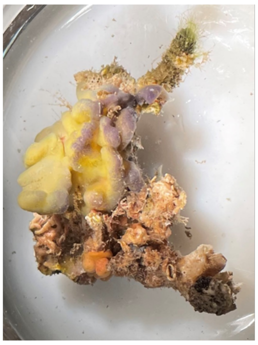

MAI_12

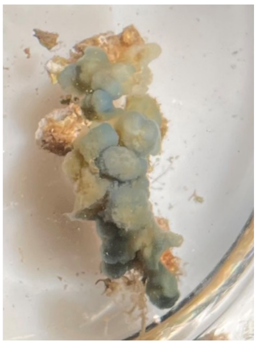

MAI_13

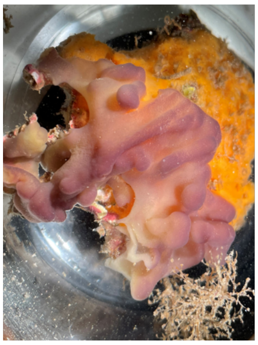

MAI_14

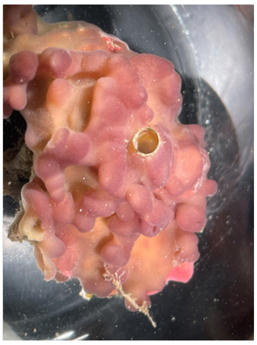

MAI_15

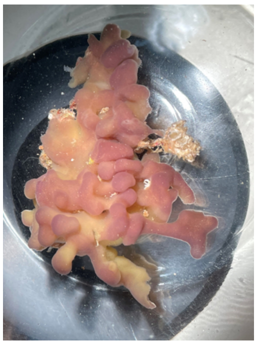

MAI_16

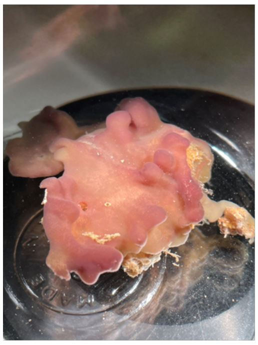

MAI_17

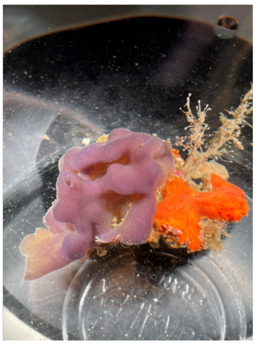

MAI_18

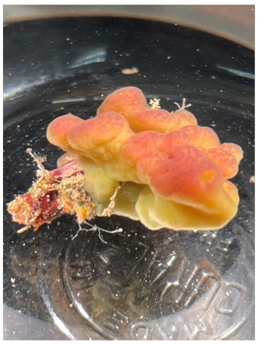

MAI_19

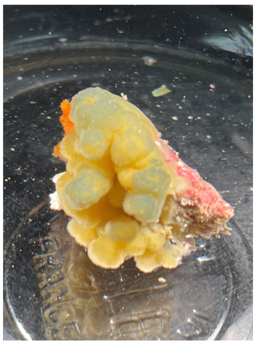

MAI_20

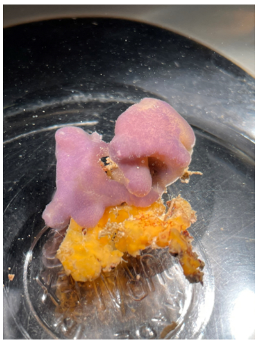

MAI_21

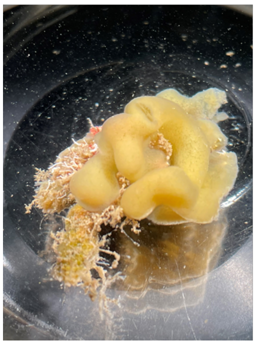

MAI_22

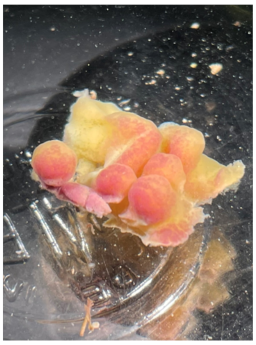

MAI_23

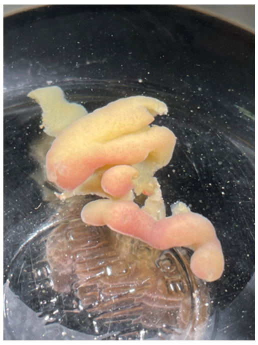

MAI_24

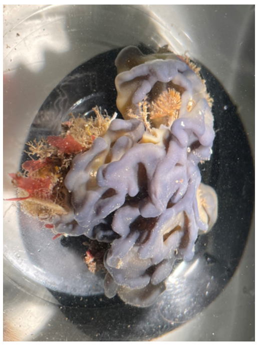

MAI_25

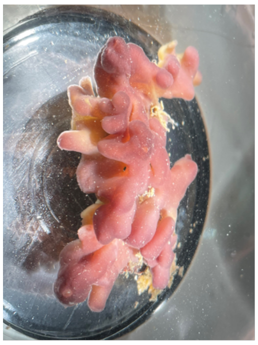

MAI_26

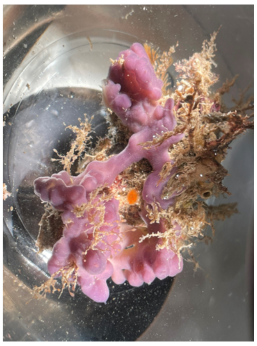

MAI_27

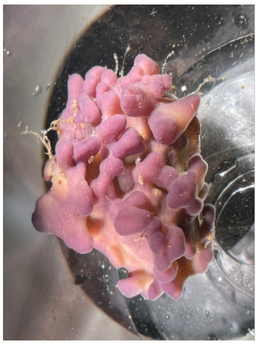

MAI_28

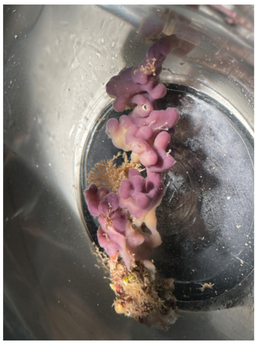

MAI_29

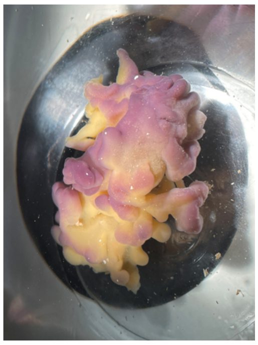

MAI_30

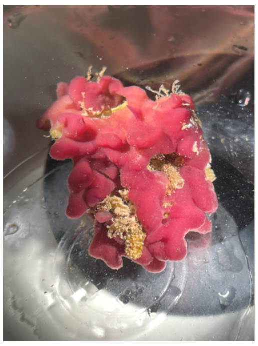

MAI_31

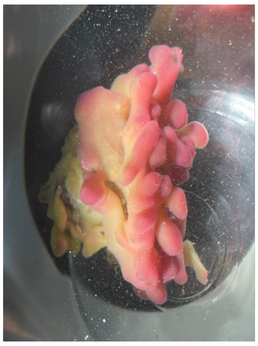

MAI_32

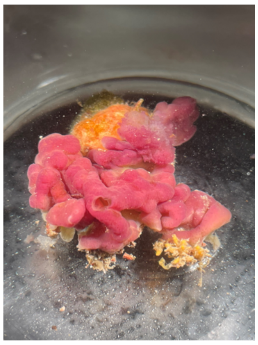

MAI_33

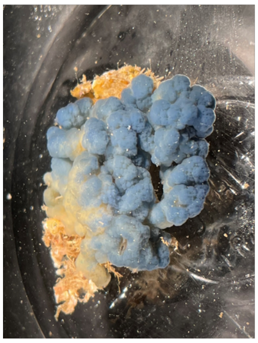

MAI_34

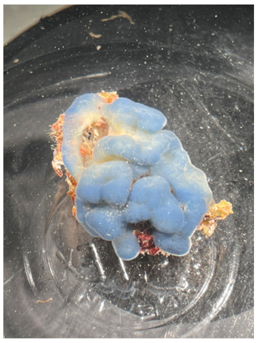

MAI_35

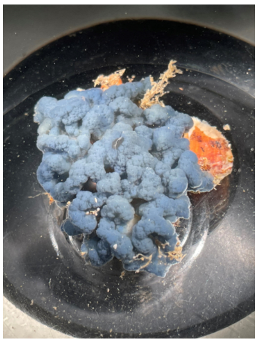

MAI_36

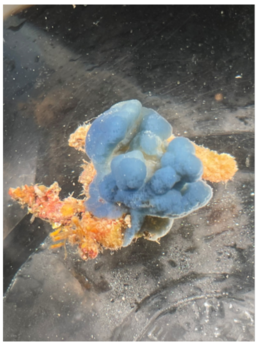

MAI_37

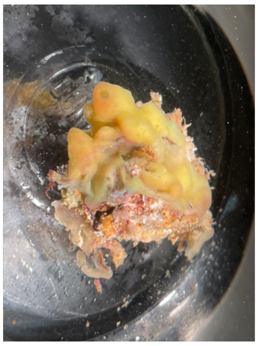

MAI_38

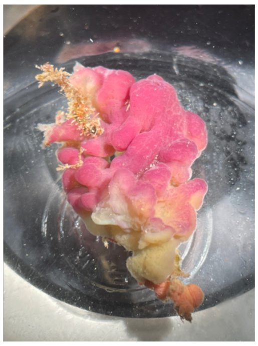

MAI_39

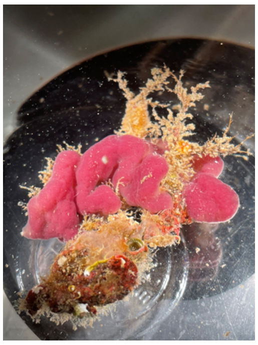

MAI_40

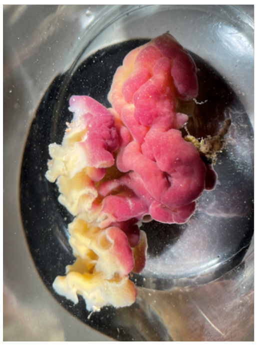

MAI_41

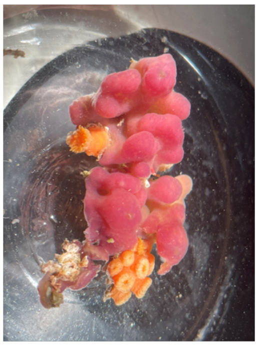

MAI_42

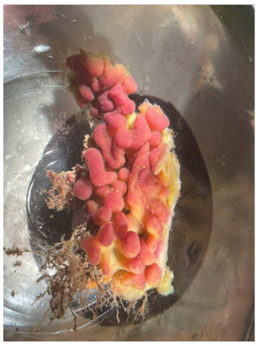

MAI_43

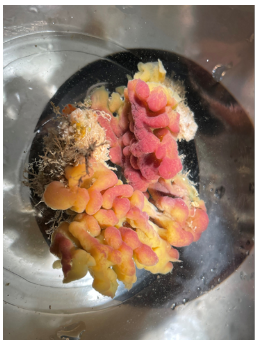

MAI_44

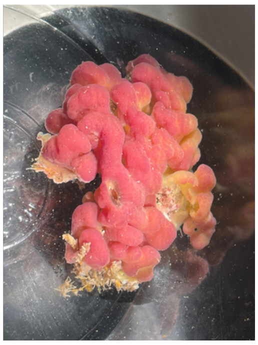

MAI_45

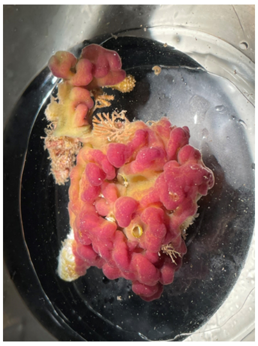

MAI_46

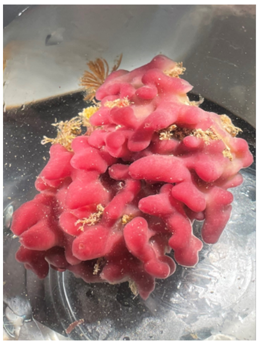

MAI_47

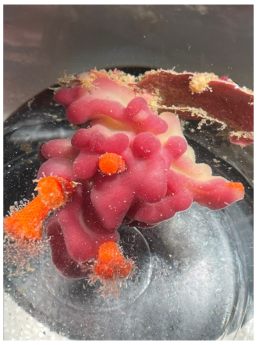

MAI_48

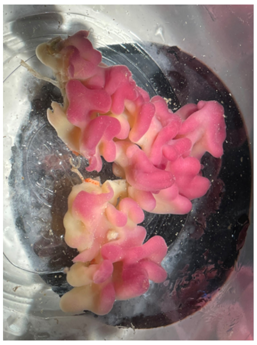

MAI_49

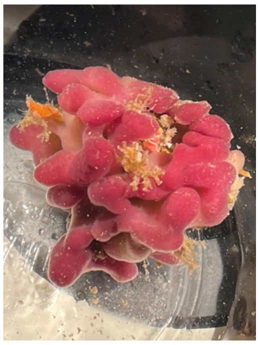

MAI_50

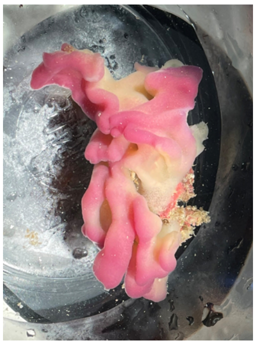

MAI_51

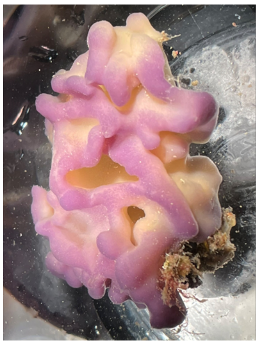

MAI_52

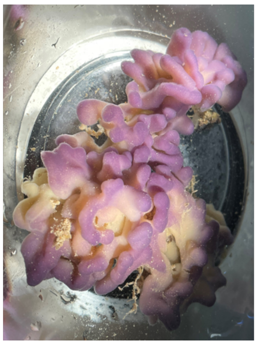

MAI_53

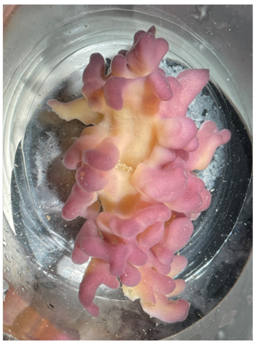

MAI_54

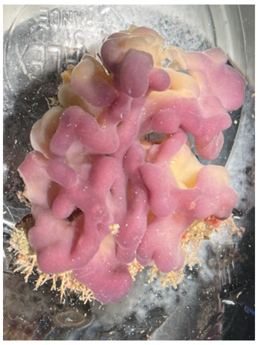

MAI_55

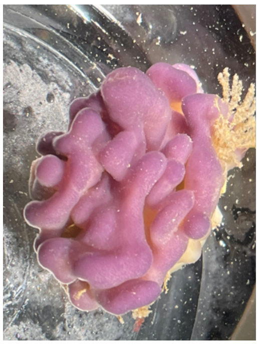

MAI_56

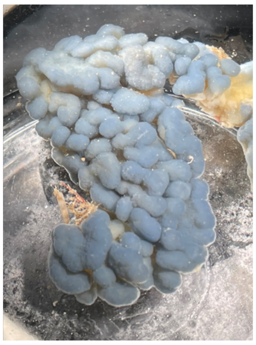

MAI_57

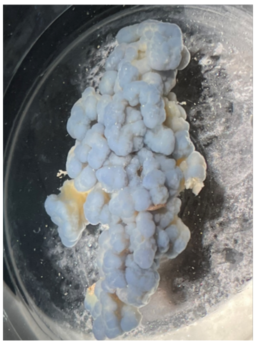

MAI_58

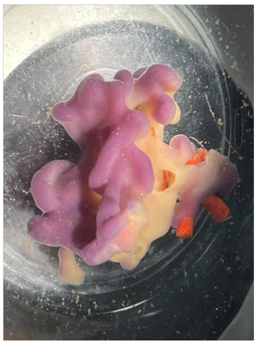

MAI_59

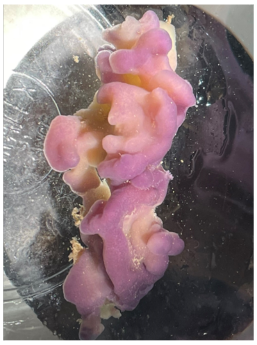

MAI_60

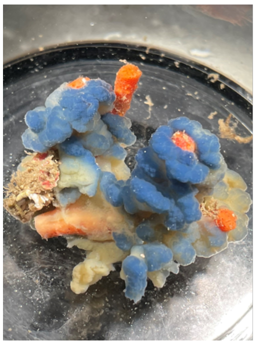

MAI_61

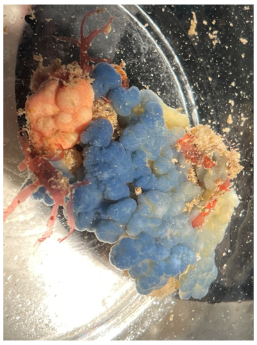

MAI_62

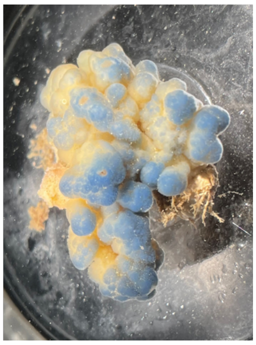

MAI_63

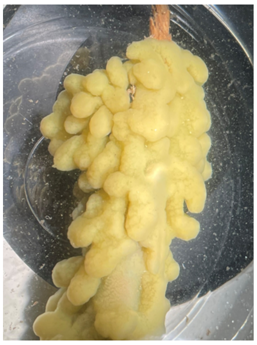

MAI_64

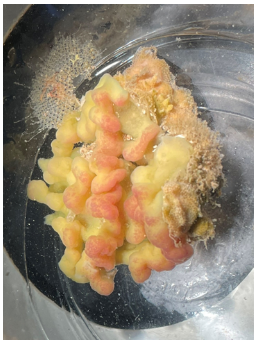

MAI_65

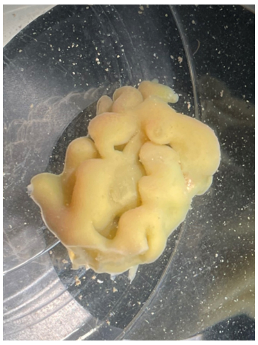

MAI_66

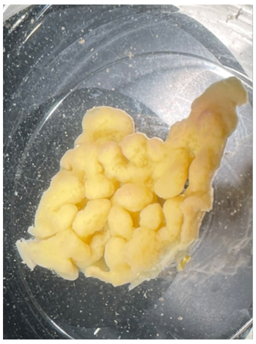

MAI_67

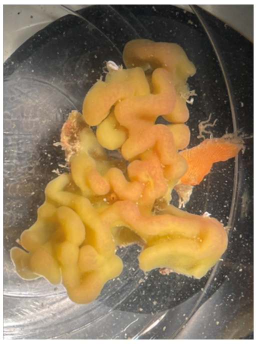

MAI_68

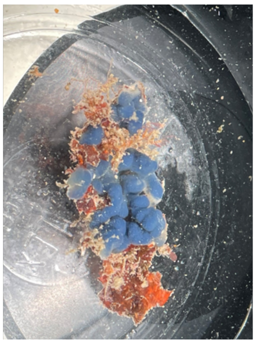

MAI_69

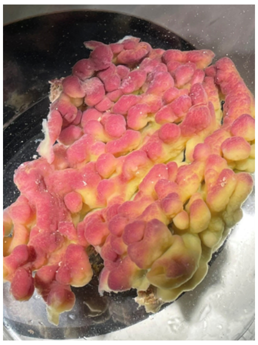

MAI_70

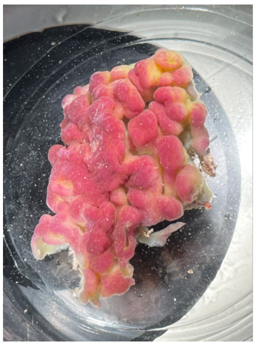

MAI_71

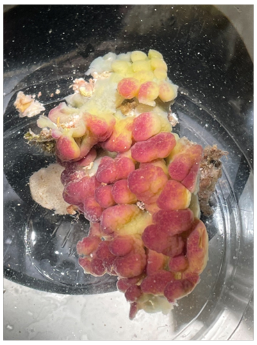

MAI_72

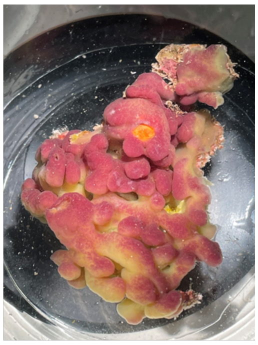

MAI_73

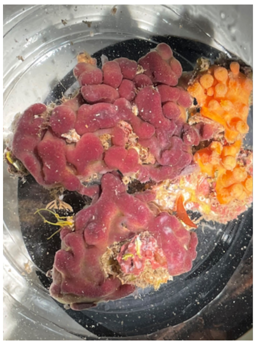

MAI_74

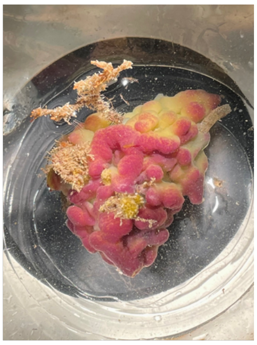

MAI_75
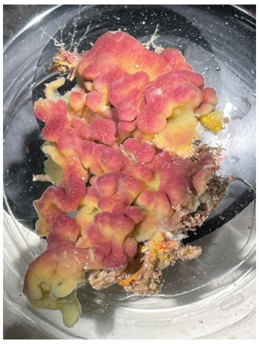
MAI_76
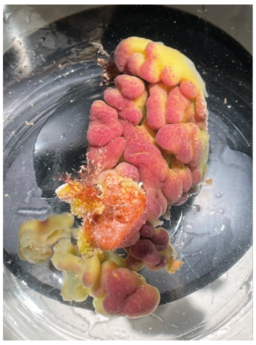
MAI_77
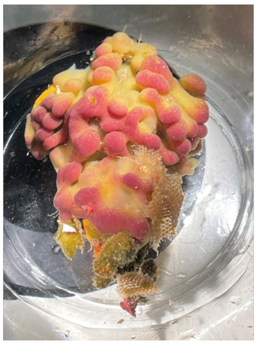
MAI_78
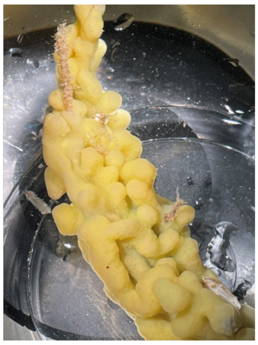
MAI_79
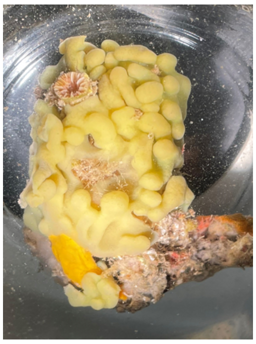
MAI_80
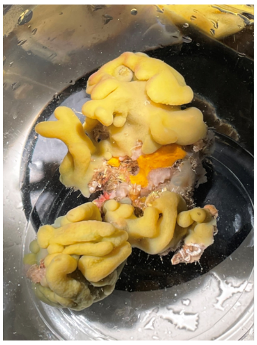
MAI_81
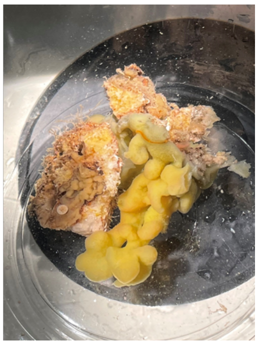
MAI_82
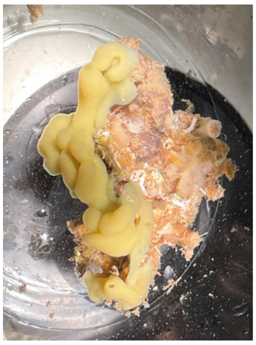
MAI_83
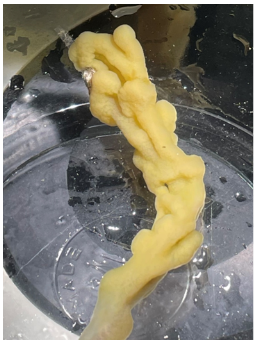
MAI_84
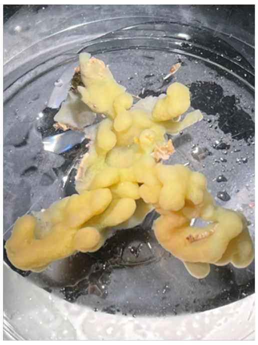
MAI_85
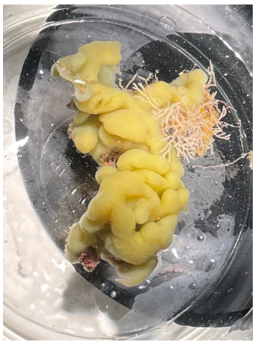
MAI_86
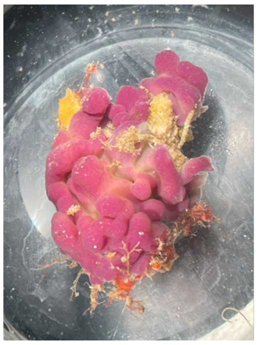
MAI_87
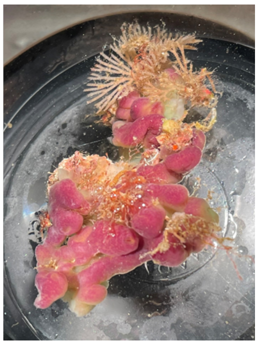
MAI_88
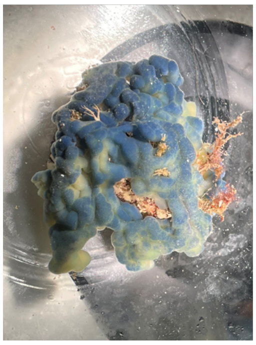
MAI_89
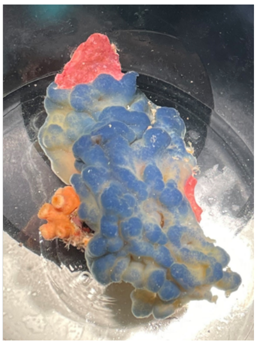
MAI_90
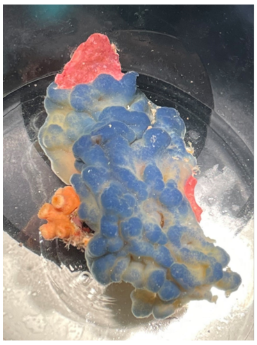
MAI_91
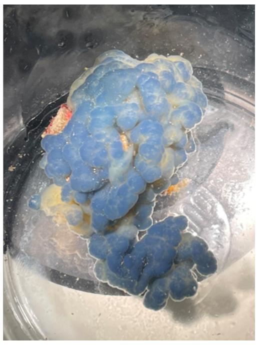
MAI_92
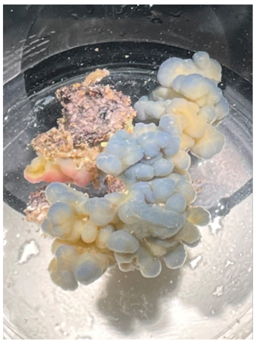
MAI_93
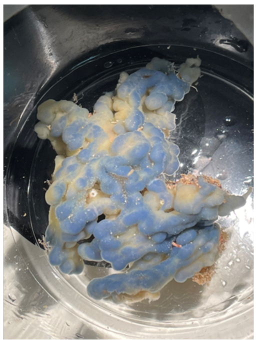
MAI_94
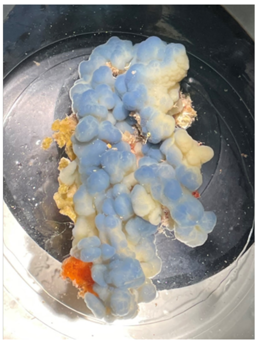
MAI_95
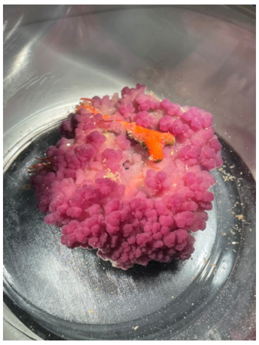
MEJ_01
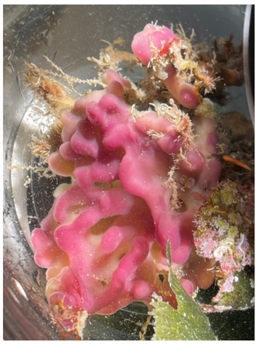
MEJ_02
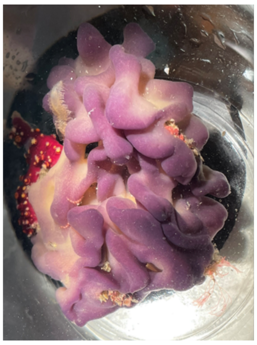
MEJ_03
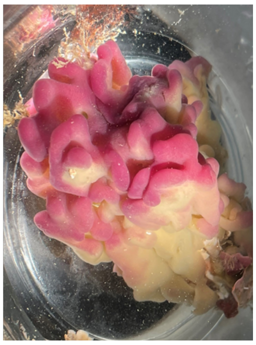
MEJ_04
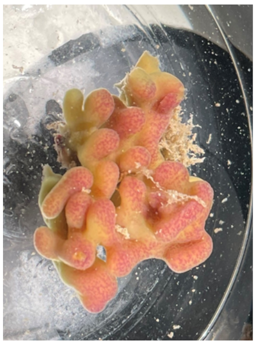

MEJ_05

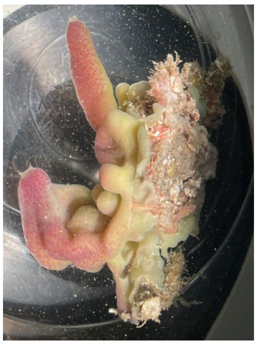

MEJ_06

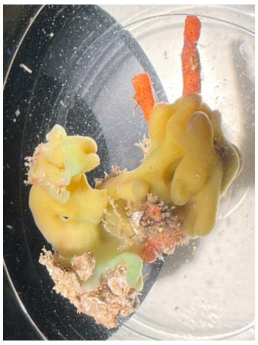

MEJ_07

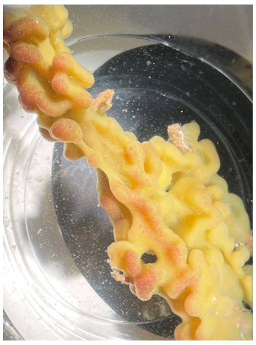

MEJ_08

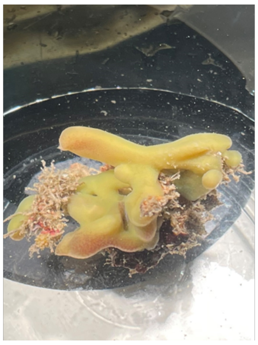

MEJ_09

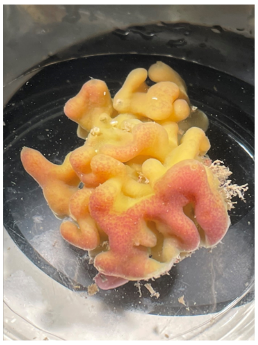

MEJ_10

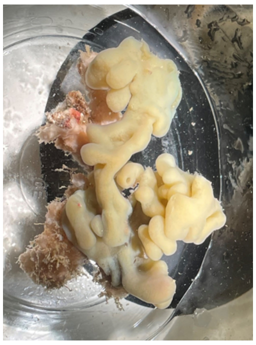

MEJ_11

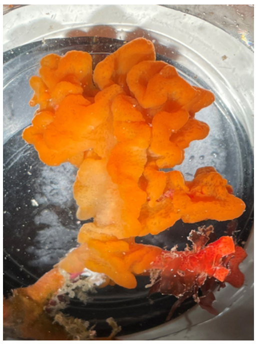

MEJ_12

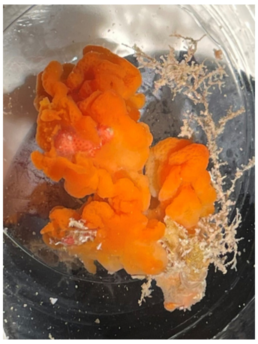

MUG_01

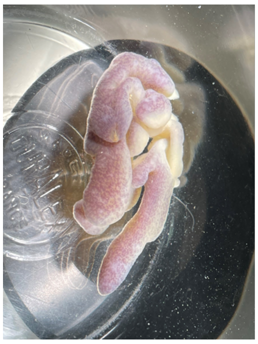

MUG_02

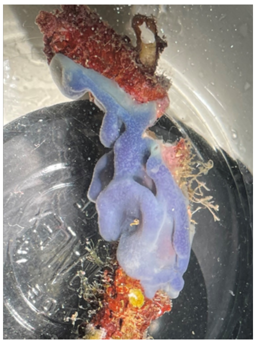

MUG_03

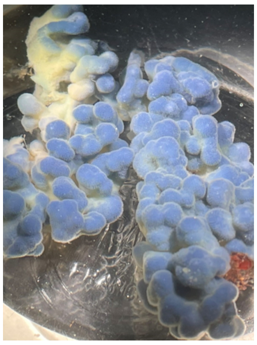

MUG_04

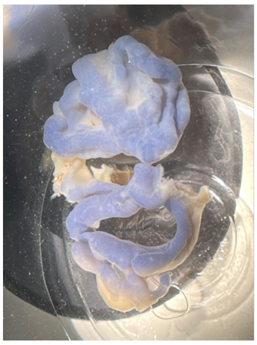

MUG_05

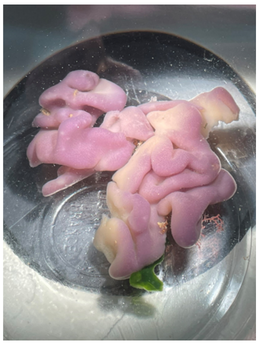

MUG_06

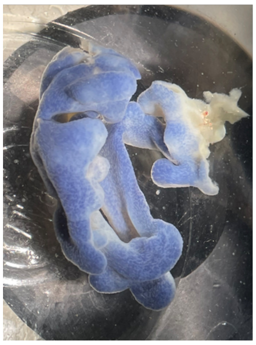

MUG_07

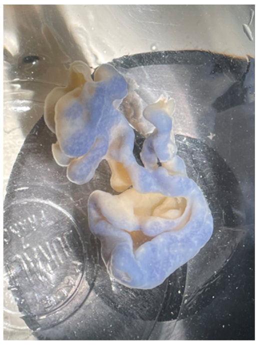

MEG_08

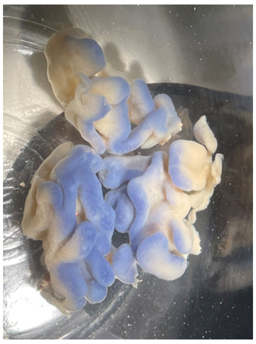

MEG_09

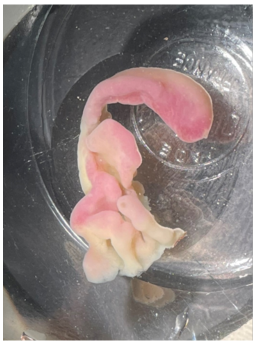

MUG_10

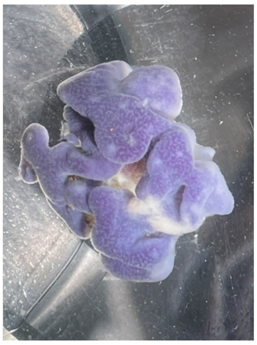

TIB_01

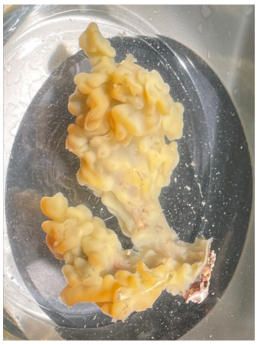

TIB_02

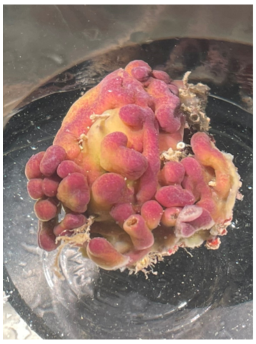

TIB_03

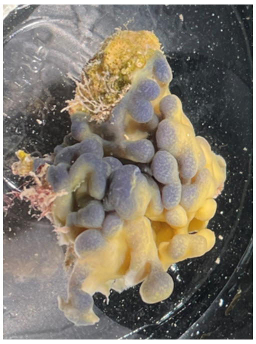

TIB_04

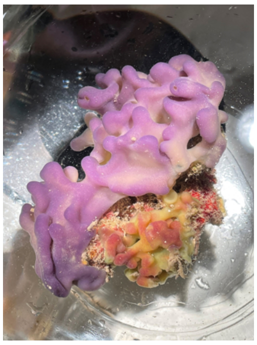

TIB_05

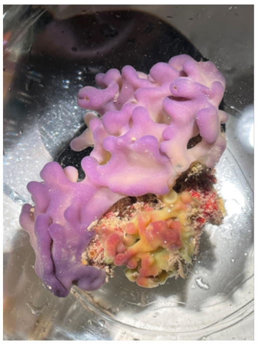

TIB_06

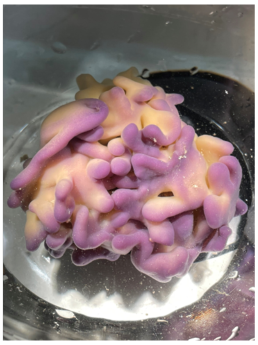

TIB_07

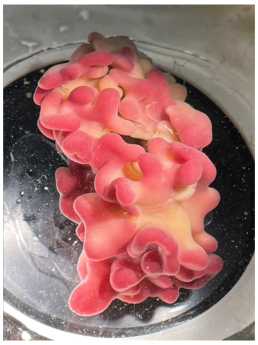

TIB_08
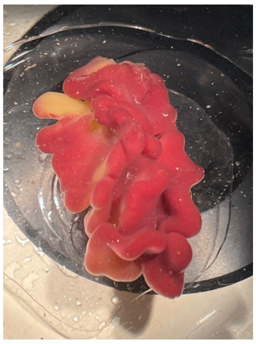
TIB_09
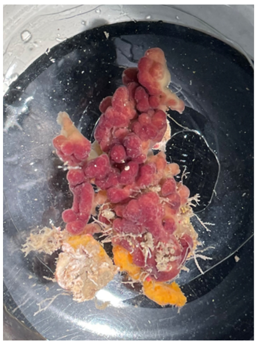
TIB_10
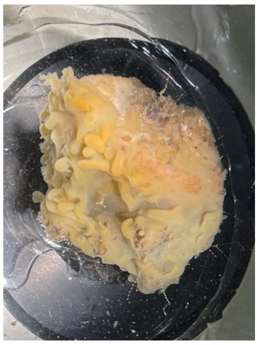
TIB_11
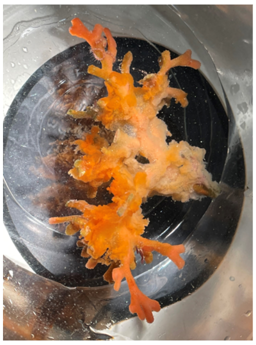
TIB_12
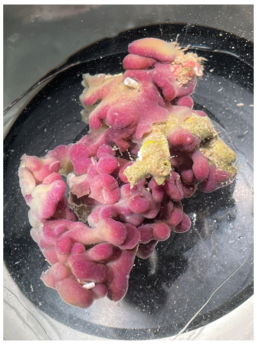
TIB_13
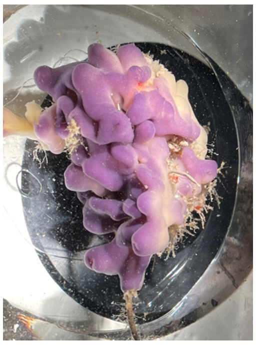
TIB_14
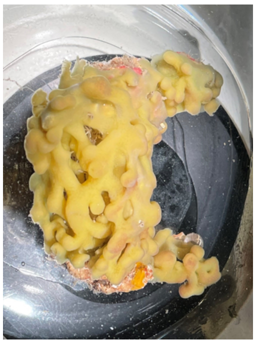
TIB_15
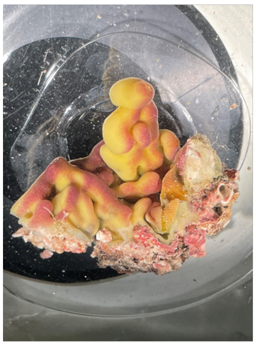
TIB_16
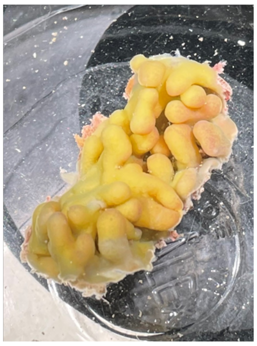
TIB_17
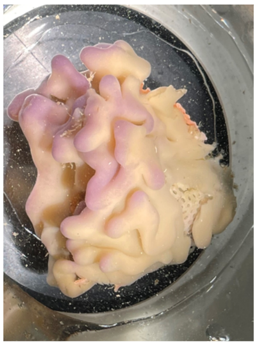
VES_01
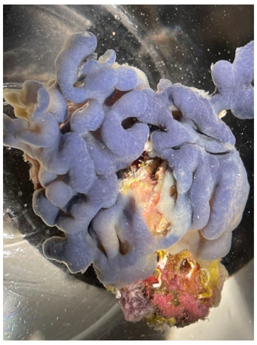
VES_02
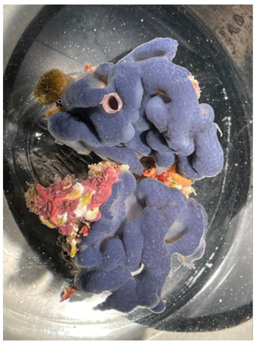
VES_03
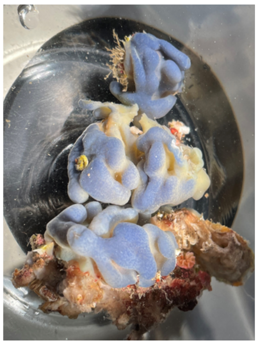
VES_04
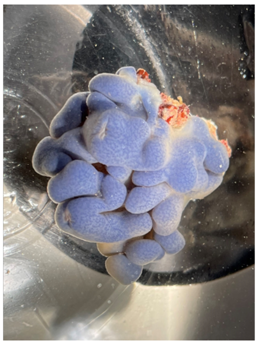
VES_05
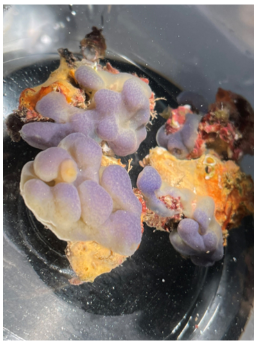
VES_06
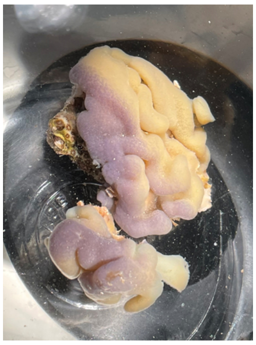
VES_07
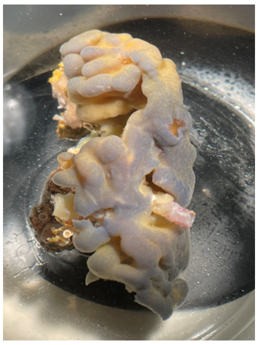
VES_08
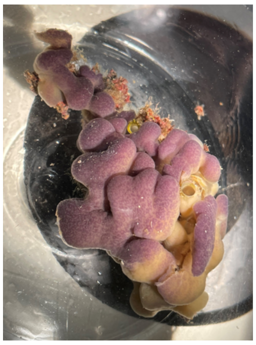
VES_09
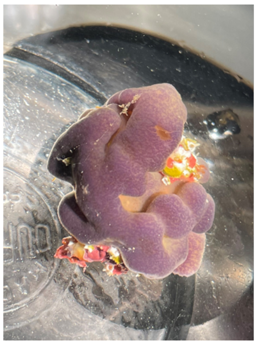
VES_10
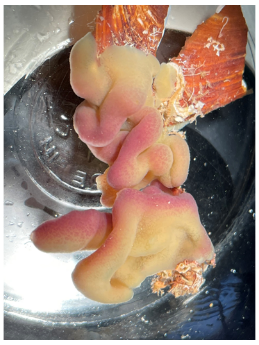
VES_11
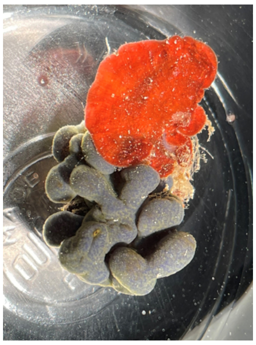
VES_12
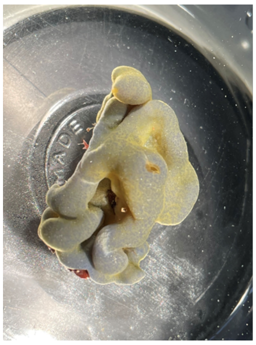
VES_13
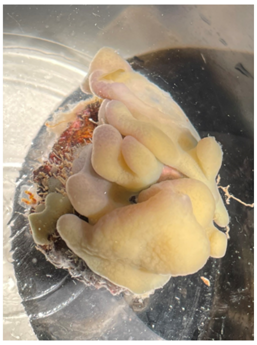
VES_14
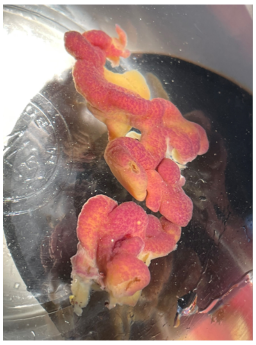
VES_15
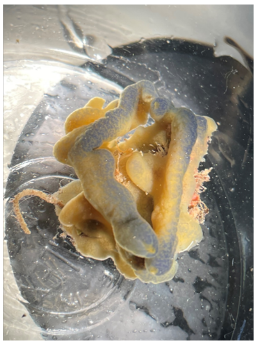

VES_16

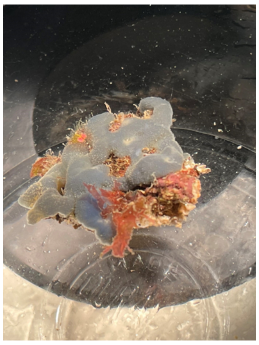

VES_17

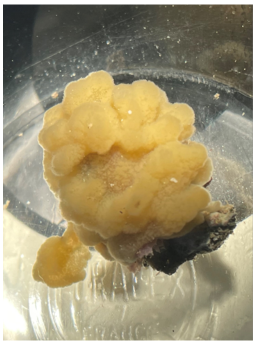

VES_18

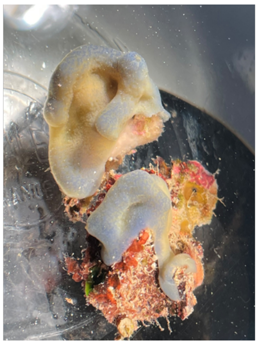

VES_19

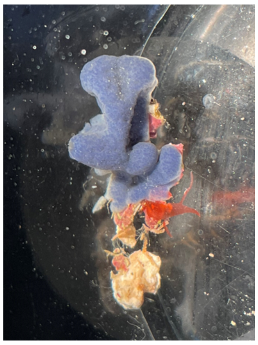

VES_20

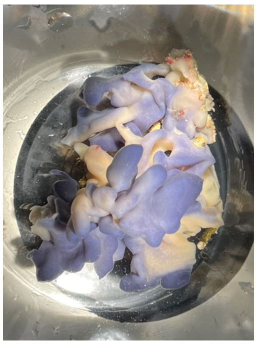

VES_21

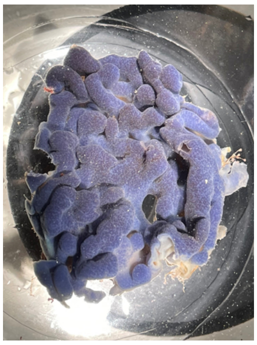

VES_22

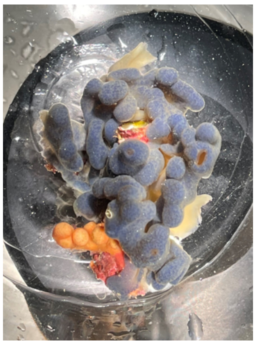

VES_23

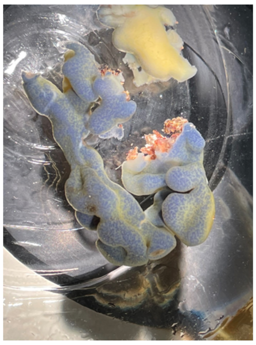

VES_24

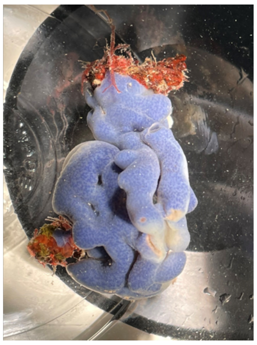

VES_25

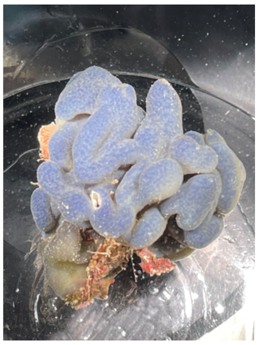

VES_26

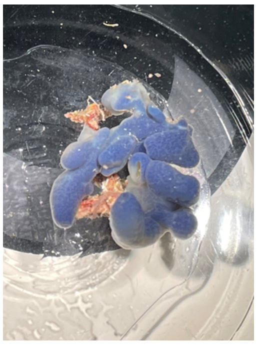

VES_27

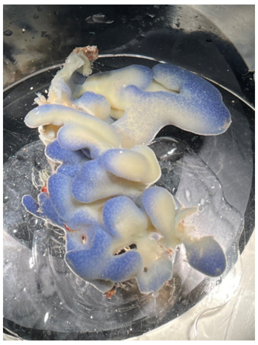

VES_28

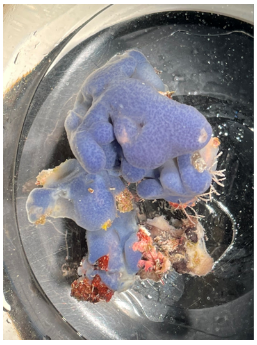

VES_29

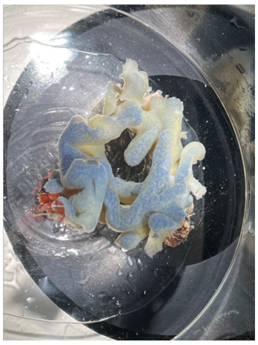

VES_30

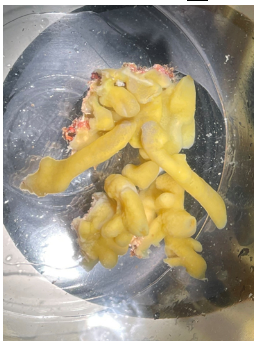

VES_31

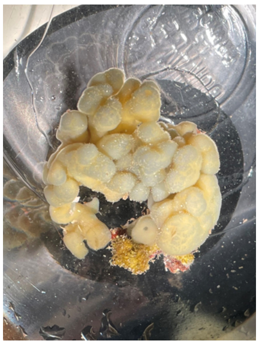

VES_32

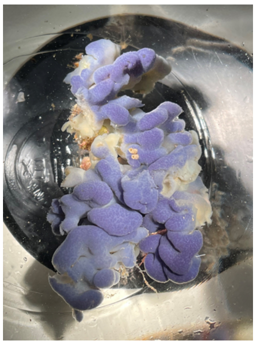

Sup. Figure 2

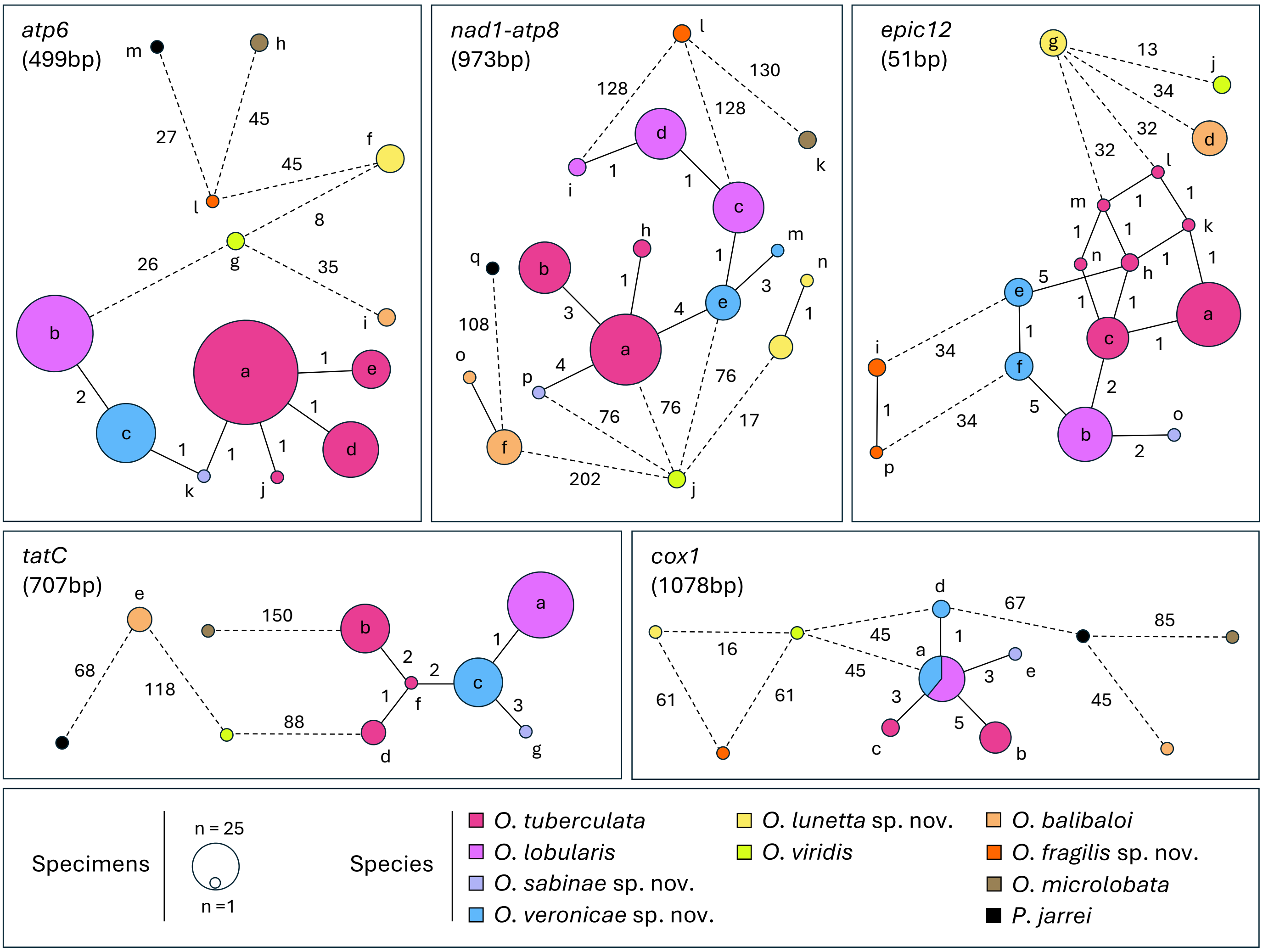

atp6
(499bp)
nad1-atp8
(973bp)
epic12
(51bp)
tatC
(707bp)
cox1
(1078bp)
Specimens
n = 25
n =1
Species
O. tuberculata
O. lobularis
O. sabinae sp. nov.
O. veronicae sp. nov.
O. lunetta sp. nov.
O. viridis
O. balibaloi
O. fragilis sp. nov.
O. microlobata
P. jarrei

Sup. Fig. 3

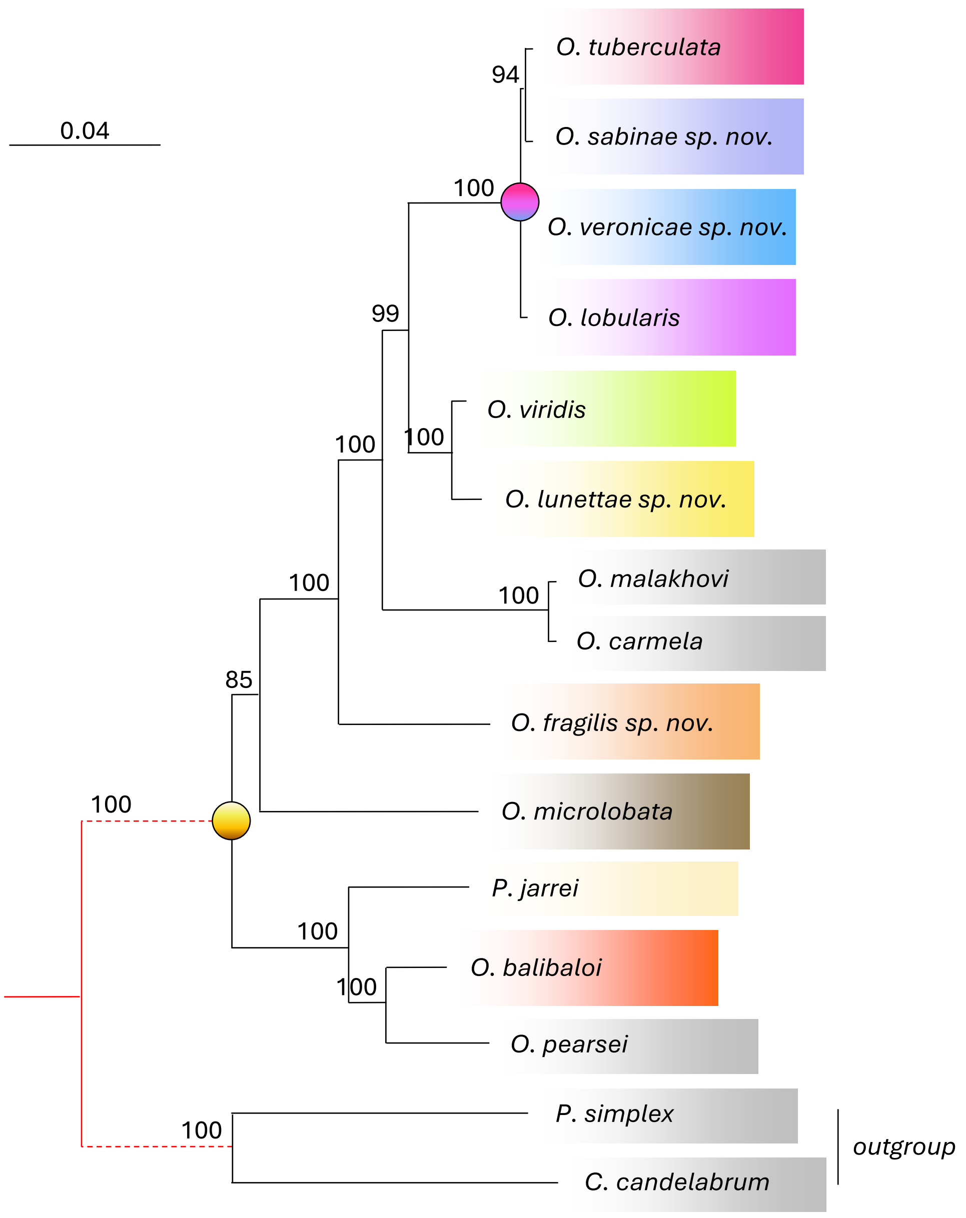

0.04
O. tuberculata
94
O. sabinae sp. nov.
100
O. veronicae sp. nov.
O. lobularis
99
O. viridis
100
100
O. lunettae sp. nov.
100
O. malakhovi
100
O. carmela
85
O. fragilis sp. nov.
100
O. microlobata
P. jarrei
100
O. balibaloi
100
O. pearsei
100
P. simplex
outgroup
C. candelabrum

Sup. Fig. 4

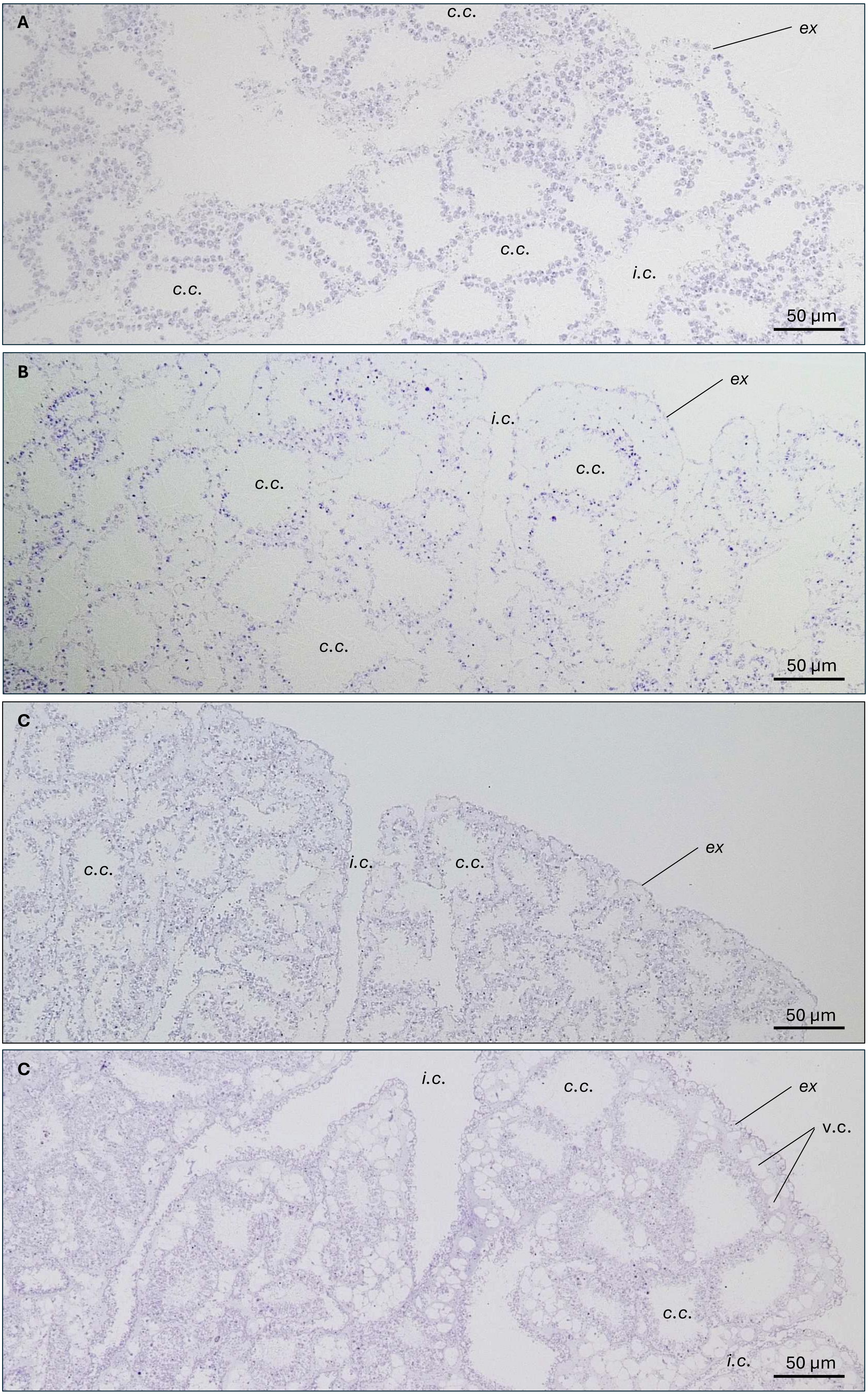
A
c.c.
ex
c.c.
i.c.
c.c.
50 µm
B
ex
i.c.
c.c.
c.c.
c.c.
50 µm
C
c.c.
i.c.
c.c.
ex
50 µm
C
i.c.
c.c.
ex
v.c.
c.c.
i.c.
50 µm

Sup. Fig. 5

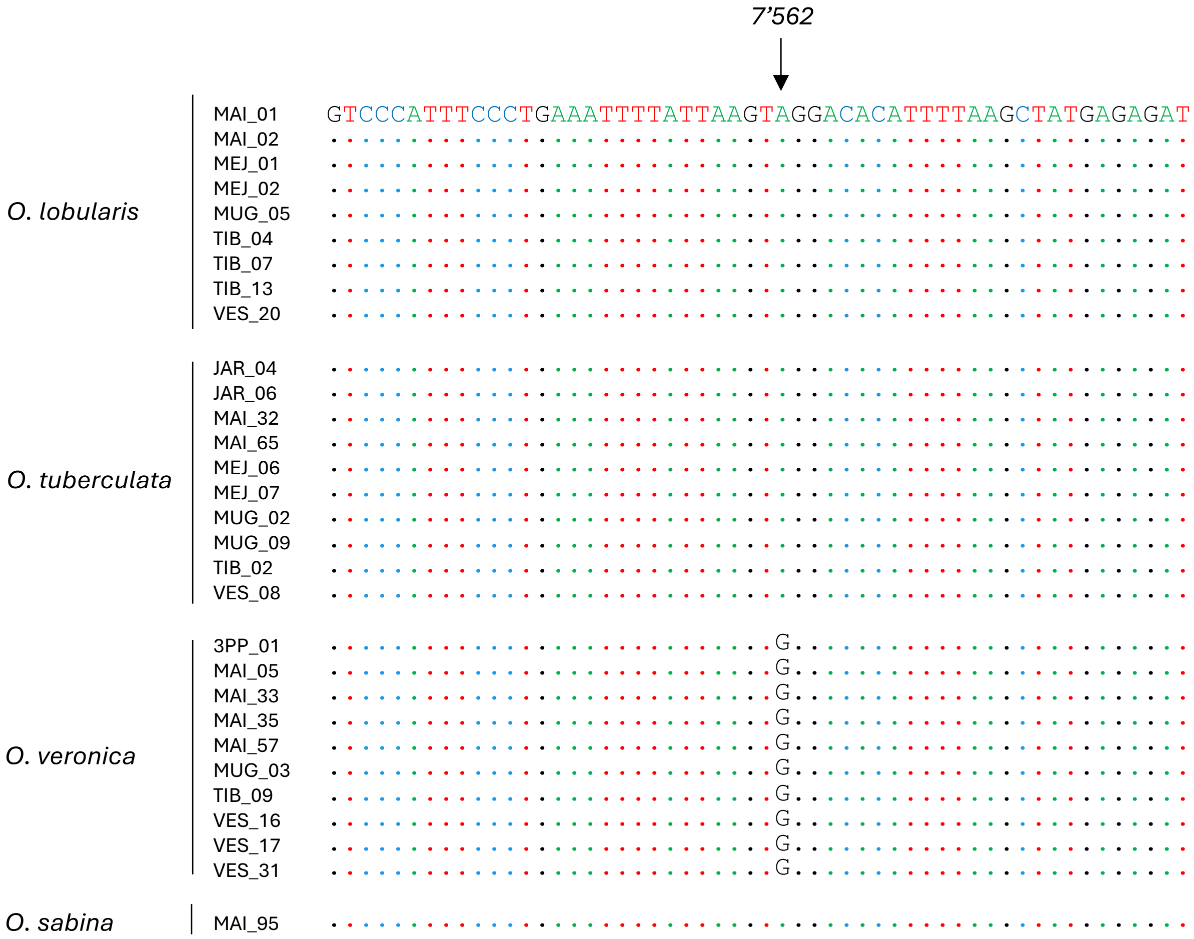
7’562
GTCCCATTTCCCTGAAATTTTATTAAGTAGGACACATTTTAAGCTATGAGAGAT
O. lobularis
MAI_01
MAI_02
MEJ_01
MEJ_02
MUG_05
TIB_04
TIB_07
TIB_13
VES_20
O. tuberculata
JAR_04
JAR_06
MAI_32
MAI_65
MEJ_06
MEJ_07
MUG_02
MUG_09
TIB_02
VES_08
O. veronica
3PP_01
MAI_05
MAI_33
MAI_35
MAI_57
MUG_03
TIB_09
VES_16
VES_17
VES_31
G
O. sabina
MAI_95
